# The biodiversity of plant-frugivore interactions: types, functions, and consequences

**DOI:** 10.1101/2025.11.14.688423

**Authors:** Pedro Jordano

## Abstract

Pairwise plant-frugivore mutualistic interactions build up into mega-diverse networks involving dozens of interacting species, being the most generalized among free-living species. These mutualisms consist of food provisioning by plants and, their counterpart, plant propagule (seeds) movement by the animals, being crucial for the natural vegetation regeneration in many ecosystems. Yet we are far from understanding which part of this enormous interaction biodiversity is needed for their maintenance. I overview the diversity of interaction modes involved in these mutualisms, the main components of the seed dispersal services, and their functional diversity. I examine how interaction richness covaries with partner species richness at different scales, resulting in variable patterns of species complementarities in terms of seed dispersal effects. The functionality of most generalized plant-frugivore mutualisms relies on complementarity of effects across a high diversity of partners, yet frequently depends on just a distinct subset of them, resulting in high functional redundancy. Two distinct aspects are relevant: 1) variable quantitative effects among species; 2) variable pairwise-interaction outcomes, between the extremes of antagonism and mutualism. Frugivory, occurring at the final stage of each plant reproductive episode, entails a large, cumulative, effect of other biotic interactions occurring at earlier stages (e.g., floral herbivory, pollination, pre-dispersal fruit damage). I examine how plant-frugivore interactions mix-up with the whole biotic interactome of a plant, using the *Prunus mahaleb* system as a case study. The effects of distinct subsets of frugivores combine with different sets of antagonistic and mutualistic partners in other interactions, yet having a lasting signal on final seed dispersal success.

**Abbreviated abstract:** Pairwise plant-frugivore mutualistic interactions aggregate into highly diverse networks crucial for seed dispersal and ecosystem regeneration, but the exact subset of interactions essential to maintain these systems remains unclear. The functionality of these mutualisms depends on complementary effects among many species, though often only a few key partners drive seed dispersal, producing both high diversity and redundancy. The outcomes of these interactions—ranging from mutualism to antagonism— are cumulative, as frugivory at the end of the reproductive cycle incorporates the influences of other biotic interactions throughout earlier plant stages, as illustrated by the *Prunus mahaleb* case study.

**Significance statement:** Pairwise plant-frugivore mutualistic interactions aggregate into highly diverse networks crucial for seed dispersal and ecosystem regeneration, but the exact subset of interactions essential to maintain these systems remains unclear. Outcomes of these interactions—ranging from mutualism to antagonism— are cumulative, as frugivory at the end of the reproductive cycle incorporates the influences of other biotic interactions throughout earlier plant stages. This study highlights the combined effects of plant-frugivore interactions within the whole plant interactome.

## Introduction

Frugivory involves a resource-based/transport mutualism supporting substantial fractions of the terrestrial biodiversity and biomass on Earth, especially in the tropics. Animal-mediated seed dispersal mutualisms depend on interaction outcomes serviced by multiple species (Corlett, 2021), so that studies focusing on pairwise interactions in isolation underestimate the levels of biodiversity required to maintain the multifunctional networks that support such plant dispersal and regeneration services. Documenting the richness, diversity, and function of these interactions requires combining approaches, such as direct and passive sampling (e.g., direct census, camera traps) and last-generation sequencing (DNA barcoding and/or metabarcoding) (Evans et al., 2016; Quintero et al., 2022) allowing the identification of species-specific contributions and estimating both demographic and genetic outcomes.

Recent work on the use of complex network theory for the analysis of plant-animal mutualisms has generated a tremendous advance in understanding how diversified interactions among free-living species evolve and coevolve (Thompson, 2009). These interactions of animals and plants take place in nature in the form of interspecific encounters, frequently involving two individuals of otherwise unrelated species. Most emphasis to date, however, has been on interactions at the species level, i.e., monitoring the patterns of mutual dependence among species in natural communities. This is just an approximation to the actual complexity of these interactions, as most often the agents involved are individuals, not species (Guimarães, 2020). This implicit aggregation of individuals into species-level defined nodes may limit our understanding of how these complex networks build up: we might fail to identify the minimum building elements (individuals, species) that are essential to form the structural bases of the mutualistic network. It is actually a sampling problem (Chacoff et al., 2012; Jordano, 2016), which is dependent on the size of these interactomes: how many species are involved in a given habitat, area or locality to support the forest regeneration system? How many distinct functions do the outcomes of their mutualistic interactions entail, and which of them are essential?

Classic work on food webs has repeatedly documented how the number of distinct pairwise interactions recorded among species within a food web scales-up as a function of increasing species richness (the total number of interacting species involved) (Sugihara et al., 1989; Bersier and Sugihara, 1997), a relation also observed in plant-animal interaction assemblages (Jordano, 1987) and, in general, in complex systems made up by component parts (Margalef and Gutiérrez, 1983). The richness of interactions a given assemblage supports thus depends on the local, *α*-biodiversity, of partners, or relates to increasing area sampled, including also some fraction of *β*-diversity (Galiana et al., 2021).

In animal-mediated, seed dispersal mutualisms, functions depend not just on the taxonomic diversity of partners but also on how their effects combine to result in effective dispersal. In these dispersal systems functions determine interaction outcomes, defined as net effects on realized fitness (final seed dispersal success). Thus, we may encounter a range of outcomes in any pairwise frugivory interaction, between the extremes of fully mutualistic (e.g., fruit consumption and seed dispersal by the so called “legitimate dispersers”) to fully antagonistic outcomes (e.g., pulp consumers and/or seed predators that may sporadically disperse seeds) (Levey, 1987; Snow and Snow, 1988). Such gradients between antagonistic and mutualistic outcomes are shared by many forms of interaction among free-living species (Gómez et al., 2023) and are ultimately related to among-partner variation in interaction outcomes. For example, Simmons et al. (2018) reported that non-mutualistic interactions with pulp peckers and seed predators occurred in seven plant-frugivore assemblages throughout Europe, accounting for 21%-48% of all interactions and 6%-24% of total interaction frequency. Increasing diversity of partners does not invariably mean increasing diversity of functional effects due to complementary interaction outcomes (e.g., McConkey and Brockelman, 2011; Schleuning et al., 2014; Rother et al., 2016), and these functional effects quickly saturate as partner diversity increases (Donoso et al., 2017; García et al., 2018). We might expect tropical frugivore assemblages to exhibit higher functional diversity (most likely associated with lower redundancy of functions) than non-tropical assemblages, frequently composed of more phylogenetically homogeneous sets of partners (Moermond and Denslow, 1985). An additional aspect to consider is that quantitative or qualitative effects are unevenly distributed among partner species. Consider for instance seed dispersal of *Prunus mahaleb* by frugivorous animals (Jordano and Schupp, 2000). Up to 75% of the feeding records are contributed by just five frugivore species among a recorded total of 38 species; the rest of the seed disperser coterie just accounts, per species, for <5% of the feeding visits to trees. This high dominance of the seed dispersal services provided by diversified frugivore assemblages visiting fruiting trees appears generalized, being consistently reported for different biomes (Quintero et al., 2025). Thus, marked inequalities in dispersal outcomes serviced by different frugivore species may limit the complementarity of these functional effects even in highly diversified assemblages. Yet it remains unexplored how functional diversity may be reduced de facto when properly weighting by interaction frequency and outcome (see e.g., Rother et al., 2016).

Frugivory involves an interaction that typically occurs at the end of each reproductive episode in fleshy-fruited plant species. Consider that frugivory on unripe fruits almost always involves seed predation and loss for the plants (Janzen, 1983). Thus, it is an interaction that, from the plant perspective, operates on the cumulative effects of myriads of previous biotic interactions occurring throughout the successive reproductive phenophases of the plants, from flower buds to ripe fruits (Harper, 1977); thus, also capturing effects of foliar herbivores, pollinators, mycorrhizae, browsers, etc. Therefore, the biodiversity of interactions with animal frugivores represents a limited, somehow reduced, subset of the biotic interactions occurring in a given reproductive episode of the plants. Recent approaches to assess multiple, distinct, types of interaction forms within complex interaction networks use multilayer networks, allowing a joint representation of the different interactions and its analysis (e.g., Pilosof et al., 2017; Hutchinson et al., 2019; Frydman et al., 2023; Hervías-Parejo et al., 2024). These multilayer structures allow the partitioning of the biodiversity of interactions among the distinct sets of partners, i.e., how each group of partners contributes to the total biodiversity of the multilayer structure.

A special type of multilayer structures are the multiplex networks (Kivelä et al., 2014; De Domenico et al., 2014; De Domenico, 2022; Bianconi, 2021), where a subset of nodes is present in any type of interaction (i.e., in any of the layers) so that interlayer links represent effects of the interaction outcomes in one layer on the status of nodes in successive layers. Multiplexity is ideal to asses, for instance, the successive interactions involved along demographic cycles with transitions across distinct stages where, e.g., individual plants in a population (nodes present in different layers of interaction) interact with specific partners (herbivores, pollinators, frugivores) which are present just in their respective layer of each interaction type. In these multiplex networks individual plants are linked through successive layers due to the fitness consequences of their interactions at each layer (Fig. 1) (Jordano, 2024, 2026); these cumulative fitness effects can be mapped in the interlayer links that connect the layers. Each layer itself maps the interactions of these plant nodes with the partner species interacting in each layer. Individual-based networks (Quintero et al., 2025) are the main elements included in each layer (Fig. 1): the plant nodes appear repeatedly in each layer because they illustrate the individuals that interact with distinct subsets of animal and fungi taxa in different types of interactions. The added value provided by such approaches is just unveiling the full potential of intraspecific variation in animal-mediated seed dispersal (Snell et al., 2019). The type of multiplexity concerns how the layers are coupled functionally, that is, how the function of one layer affects that of another (and vice versa) (Lee et al., 2015). In our case example such interlayer coupling would be defined in terms of fitness effects acting in a connective manner through sequential effects at different demographic stages (Fig. 1) (Jordano, 2024, 2026).

**Figure 1:**
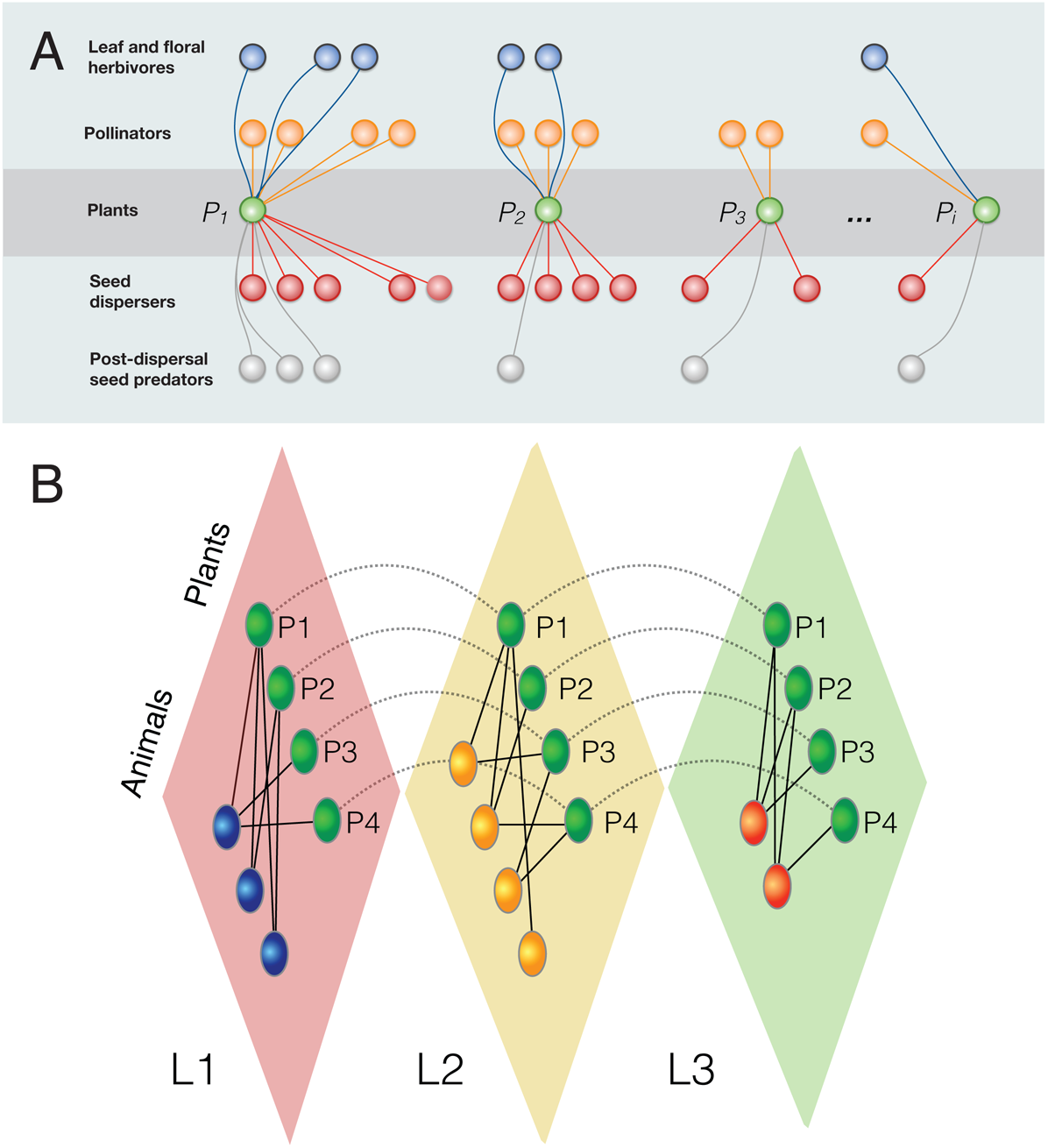
Multiplexed ecological interactions in the life-history demographic cycles of plants (Harper, 1977). A, Different individual trees *P*_1_…*P*_*i*_ with their respective interactomes (sets of partner species in different types of interaction, e.g., herbivory, pollination, seed dispersal, mycorrhizae). Outcomes of these interactions involve fitness consequences, which are cumulative across the succesive stages of the reproductive cycle illustrated by different layers (*L*_1_…*L*_3_). Multiplexity is illustrated in B by interlayer links (dashed)representing the fitness consequences of each interaction type on the status of the plant nodes in successive layers. The same plant node (state nodes) is present in each layer, while each partner taxa is only present in its layer. The interlayer links represent the fitness consequences of each interaction type on the status of the plant nodes in successive layers.

My objectives here are to provide a broad overview of plant-frugivore interaction biodiversity-i.e., how many distinct interactions?-emphasizing not just the diversity in quantitative terms of interaction frequencies, but also in terms of assemblage complementarity/redundancy. Then, I examine how plant-frugivore interactions are assembled within larger and more complex webs of interaction with other biotic partners like herbivores, fungi, and pollinators to test for the distinct signals of plant-frugivore interaction outcomes on realized fitness (dispersed seeds).

## Material and Methods

### Interaction richness and diversity

I compiled data on number of species (plants and animal frugivores) and number of interactions included in different compilations (e.g., Jordano, 1987; Fricke and Svenning, 2020; Martins et al., 2022; Dehling et al., 2021; Mendes et al., 2024) and repositories (WebofLife [https://www.web-of-life.es]) of plant-frugivore interactions, and with added studies (see Suppl. Mat. Table S1). Most datasets refer to local assemblages (i.e., studies in a single locality or multiple localities within reduced regions) of fleshy-fruited plant species and animal frugivores, yet others span over larger regional or continental or sub-continental areas (e.g., Mendes et al., 2024) or include interactions of fruit consumption/potential dispersal of non-fleshy-fruited plant species. This totalled *N*= 168 datasets with information on both species and interaction richness. Linear regression of *I*, the total number of distinct pairwise interactions recorded, on *S*, species richness of plants and animals combined, were estimated with R package lme4 (Bates et al., 2015). I used either weighted or non-weighted, binary, adjacency matrices (Suppl. Mat. Table S1).

### Interaction richness and functional traits

To assess functional diversity patterns in relation to species richness in frugivore assemblages I resorted to individual-based interaction networks (Quintero et al., 2025) that represent the interactions between individual plants in local populations and the local frugivore species. The rationale is to explore how the functional diversity of partners for plants within a specific population increases as the number of partner species in the mutualistic assemblage increases. I used just a subsample of the individual-based networks listed in Suppl. Mat. Table S1, specifically those of *Prunus mahaleb* (L.) (Rosaceae) and *Euterpe edulis* (Mart.) (Arecaceae) (Jordano and Schupp, 2000; Galetti et al., 2013).

I evaluated the functional diversity *FD* of the frugivore assemblages for both species, estimating several functional diversity indexes with the *R* package *FD* (Lavorel et al., 2008; Laliberté and Legendre, 2010). Specifically, I examined *FD* variation using functional richness, *FRic*, functional evenness (*FEve*), and functional divergence (*FDiv*), as well as functional dispersion (*FDis*), and Rao’s quadratic entropy (*Q*) (Laliberté and Legendre, 2010). I used several species traits to assess functional diversity of the two frugivore assemblages [trait data coming from (Galetti et al., 2013; Jordano and Schupp, 2000; Bovo et al., 2018; Jordano et al., 2025)], including, body mass, gape width (measured at the mouth commissures), visitation rate to the trees (no. visits per 10h of observation), percentage of fruits swallowed (vs. pecked), seed dispersal effectiveness (both per visit and total; Schupp.etal:2017:EL).

All indices were weighted by the relative visitation frequency of each frugivore species to the plants. *FD* indexes are evaluated based on two distinct datasets with species-specific traits, one with ecomorphological variables and another with abundance values. For *P. mahaleb* abundance data were derived from *N*= 43 transect censuses of 66.4 km total length and totalling 4218 birds recorded. For *E. edulis*, abundance data refer to point-count-abundance data *IPA* from transect censuses in non-defaunated areas of SE Brazil Mata Atlantica (Galetti et al., 2013). Visitation data (no. vists per 10h) were obtained by direct focal watches at fruiting trees for 336 h (16 tree-days per two seasons) in *P. mahaleb* and 2326 h in *E. edulis*. As a weighting factor for ecomorphological data to use with *FD* indexes estimates, I considered visitation rate (i.e., frequency of interspecific encounter) to the trees rather than local abundance. Both are significantly correlated across species (*r*= 0.5953, *P* < 0.0001, *N*= 26; *r*= 0.9260, *P* << 0.0001, *N*= 24; for *E. edulis* and *P. mahaleb*, respectively, Fig. Suppl. Mat. 1).

The species-specific contribution to *FD, contrib*_*FD*_, was computed as the product of the species’ visitation rate (vis.10h) and total *FD* value for the plant, divided by the total number of visits of all frugivore species (Basile, 2022). *FRic*, functional richness, can be interpreted in our context as the distinctness of functional roles among species in a frugivore assemblage. Increasing functional richness likely means increasing the diversity of potential outcomes of the functional seed dispersal service provided by a specific frugivore assemblage. For example, increased spread of the seed shadows among multiple microhabitat types, or along a broader span of seed dispersal distances. *FDis* is a multidimensional index of functional dispersion, i.e., the mean distance of individual species to the centroid of all species in the community. It can be interpreted as a measure of the functional spread of the set of species, i.e., a measure of the functional heterogeneity of the species. Increasing functional dispersion can be interpreted in our context as increased complementarity of the seed dispersal services (see e.g., Schleuning et al., 2014; Donoso et al., 2020); thus, the addition of functionally “redundant” species would not necessarily result in increased *FDis* despite some increase in *FRic*.

For the two frugivore assemblages I analysed how functional richness *FRic* and functional dispersion *FDis* vary with increasing species richness of partners. I generated simulated assemblages by bootstrap-resampling the functional data set (without replacement, *N*= 10 resamplings), yielding assemblages increasing in number of frugivore species from 5 species to the total number of species recorded in each assemblage (28 species for *Euterpe edulis*, 26 species for *P. mahaleb*). I choose 5 species as a minimum as no convex hull can be computed when there are more dimensions (i.e., traits) than points (i.e., species) (Laliberté and Legendre, 2010).

### Case study: multiplexed plant-animal interactions in *Prunus mahaleb*

I analyse data coming from a long-term study with *Prunus mahaleb* (L.) (see Jordano, 1993; Jordano and Schupp, 2000, for details on the study system) to assess how plant-frugivore interactions are embedded within more complex networks of biotic interactions. The study species, the Saint Lucie’s or Mahoma’s cherry, is a small tree (2-10 m in height) that grows scattered at mid-elevations (1250-1900 m) in the southeastern Spanish mountains, through the Pyrenees and central and eastern Europe to Ukraine, and from Morocco through Syria to west-central Asia (Webb, 1968).

A total of *N*= 43 trees were monitored in a 25 ha area of the Sierra de Cazorla, southern Spain, during six reproductive episodes including the whole flowering and fruiting seasons (1988-1993). The study area is a Mediterranean montane forest with a rich assemblage of fleshy-fruited plants and frugivorous animals. A subset of *N*= 19 trees was monitored for herbivore (fungi, invertebrates and vertebrates) presence and interaction frequency, pollinators, flower and fruit production, frugivore visitation, and seed dispersal success. Sampling details are provided in Suppl. Mat. Briefly, I used direct counts at the individual trees, concentrated during leafing in early spring and then during flowering and fruiting. Counts involved direct censuses of 10-15 min in different parts of the tree crown (for herbivores and pollinators), and were repeated at different times of the day between mid-April and late May, encompassing the leafing and flowering phenophases of the trees (see Jordano, 1993, for details on pollinator surveys). For fungi and vertebrate herbivores, the counts of damaged leaves and branches were scored as damage levels for later analysis (see Suppl. Mat.). For animal frugivores I used direct focal watches at fruiting trees, with 10h of observation per tree and season (see Jordano, 1993, 1995; Jordano and Schupp, 2000, for details on census protocols, and estimation of flower and fruit crop sizes). Individual trees were individually tagged and geo-referenced (Jordano, 1995; García et al., 2009), recording data on tree size, flower and fruit crop size, spatial location (distance to nearest two neighbours), and fruit consumption and seed removal success (see details in Jordano, 1995, and Suppl. Mat.).

Interaction records with fungi, herbivores, pollinators, and animal frugivore taxa were summarized as adjacency matrices, one for each type of interaction. Each of these matrices had plants (individual trees) and animal/fungi taxa as nodes, with the tree nodes present in each of the three layers (state nodes corresponding to each position in the multiplex arrangement associated to the specific type of interaction considered). In contrast the animal/fungi taxa nodes were physical nodes appearing in each of the interaction type where the taxon participated (Fig. Suppl. Mat. 2). Multiplexing is thus defined as the repeated appearance of the same individual trees in separate networks depicting their interactions with different sets of partners depending on interaction type (herbivory-including also fungi, pollination, and seed dispersal). The individual tree nodes in each layer are thus linked across successive layers by means of interlayer links (see Fig. 1). Yet, given that I don’t focus on fitness effects, interlayer links were set to the same value for the sake of simplicity, just to illustrate the connections among successive interaction stages. For our purposes here interlayer link strengths were later estimated with Infomap (Edler et al., 2017; Frydman et al., 2023). Both data and code for analyses are available as Suppl. Mat. and in the GitHub repository ^7^ (https://github.com/pedroj/MS_Oikos_SI-FSD2024).

## Results and Discussion

### The biodiversity of plant-frugivore interactions

Plant-frugivore assemblages typically consist of aggregated interactions with animal partners that occur at two levels: first, individual plants of each species have individual-specific partners that characterize their individual coteries of mutualists/antagonists for seed dispersal. Second, in any given locality or site, we can aggregate those individual-based assemblages at the species level, for each plant species, and then assemble all the locally-coexisting species, resulting in species-level assemblages and the complete interaction network. Such species-level coteries have been traditionally studied as seed disperser assemblages and characterize the way a given species interacts with the suite of animal partners available.

Across assemblages at different spatial scales, from local sites at, i.e., 20-50 ha plots, to regional areas, and to biome- and continental scales, the number of distinct pairwise interactions, *I*, among sets of *P* plant species and *A* animal frugivore species, increases steadily with *S*, the total species richness of the mutualistic assemblage (*S* = *A* + *P*), (Fig. 2; *log*(*I*) = –0.1493 + 1.2632(*log*(*S*)), *Adj R*^2^ = 0.9120, *P* ≪ 0.0001, *d*.*f*. = 164). It is interesting to visualize how this increase in species richness, however, results in highly sparse interaction networks, with relatively low connectance (Jordano, 1987), a pattern also generalized in food webs (Cohen, 1978; Margalef and Gutiérrez, 1983; Briand and Cohen, 1984). The total number of interactions *I* present in a given system would be thus the result of variable combinations of *P* and *A* values, i.e., the respective richness of plants and animals in the interacting assemblages. However, plants do have a slightly higher contribution to interaction richness, as evidenced when teasing apart the effects of *P* and *A* on *I*: *I* = 0.8455 + 0.6557*P* + 0.5987*A*, with both the plant and animal effects being significant (*P* ≪ 0.001) (difference in *β* = 0.057).

**Figure 2:**
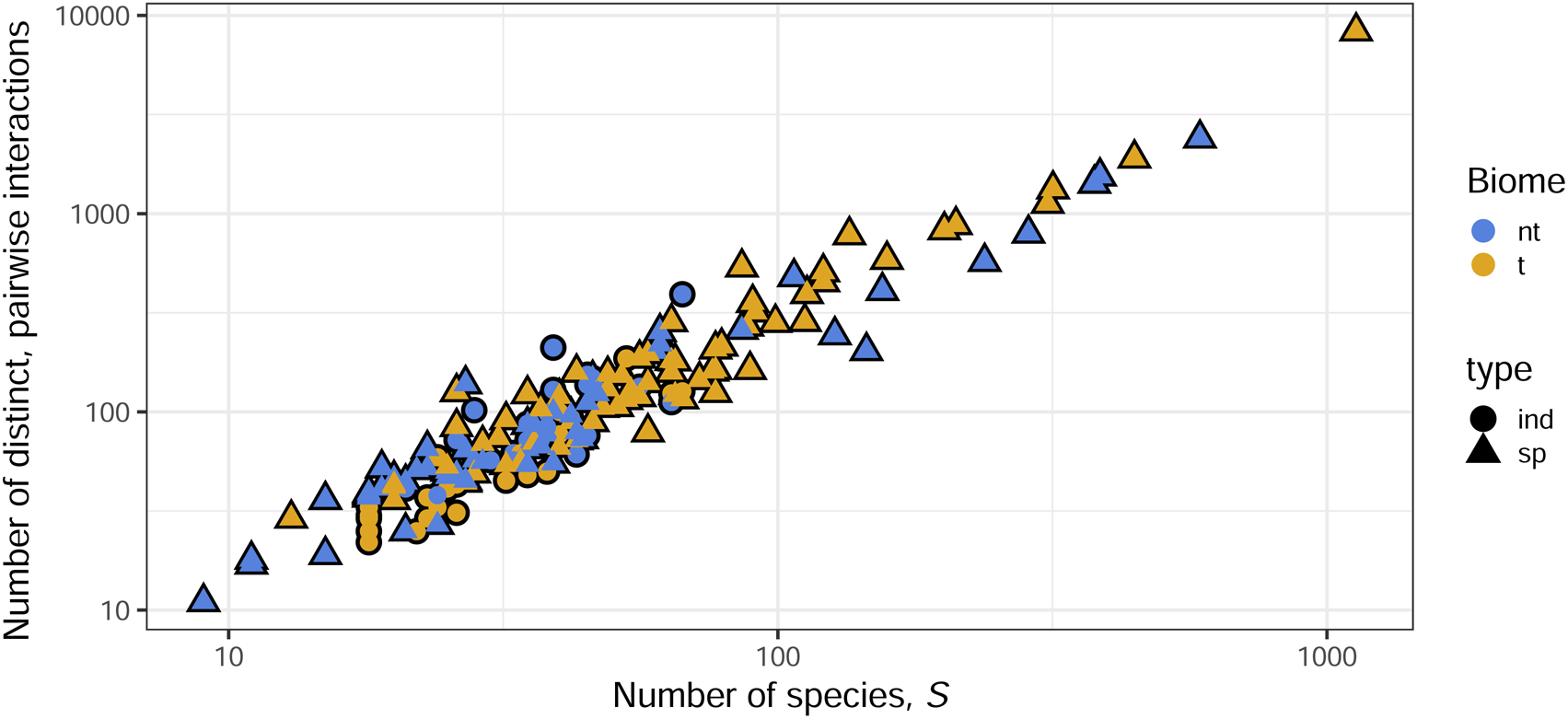
The number of distinct pairwise plant-frugivore interactions *I* in local assemblages increases linearly (log-log plot) with species richness of partners *S* = *A* + *P* (the combined species richness of plants and frugivores). Biome types: nt, non-tropical; t, tropical. Network types: ind, individual-based; sp, species-based. Data sources are listed in Suppl. Mat. Table S1.

Interestingly, the tropical and non-tropical assemblages differ in the way new interactions are added when new species are added to these networks: the slopes of the regression of *log*(*I*) on *log*(*S*) differ significantly, *b*_*trop*_ = 1.3345 ± 0.0405 and *b*_*nontrop*_ = 1.1917 ± 0.0457, respectively, with the biome effect being significant in the ANCOVA (*F* = 6.11, *P* = 0.014, *d*.*f*. = 1,163). This suggests tropical systems add almost two times more interactions per species than non-tropical systems, a pattern that has been previously reported for plant-frugivore assemblages (Jordano, 1987). Yet, due to the expected higher *S* of tropical assemblages, the resulting connectance of the interaction networks is lower in the tropics (*mean* = 0.267 ± 0.02, *n* = 91) when compared to non-tropical areas (0.332 ± 0.02, *n* = 76), as expected from the generalized decay of *C* with *S* in any interactive complex system (Margalef and Gutiérrez, 1983).

When considering individual-based interaction networks, however, we don’t expect such a steep increase in interaction richness with increasing species richness, as the number of interactions is limited by the number of individuals in each population. In addition, the heterogeneity in frugivore partner species is expected to be higher among different plant species than among different individual conspecific plants (Quintero et al., 2025). In fact, the number of interactions recorded in individual-based networks is just marginally lower than that expected from the species-level assemblages (Fig. 2), with no differences in the regression patterns. For species-based networks, *I* = –0.0036 + 1.233*S*; for individual-based networks, *I* = –1.042 + 1.498*S* (note *S* for this latter regression is the number of individual trees in the network); and the slopes of the two regressions are significantly different (*ANCOV A,F* = 4.34, *P* = 0.039). Moreover, these slopes are pretty similar to those previously obtained for food webs (Martinez, 1992) and host-parasite interactions (Dallas and Jordano, 2021), emphasizing shared scaling laws among all these trophic interactions. Other things being equal, a locally abundant tree species with a large number of individuals is expected to have a larger number of interactions than a locally rare species with few individuals, even if both species have the same number of frugivore partners. It is well documented (e.g., Jordano, 1987; Vázquez and Aizen, 2003; Vazquez et al., 2009) that local abundance is a major driver of interaction richness in these generalized mutualisms involving free-living animal frugivores and fruiting plants.

Here I’m not examining the relationship between interaction richness and area sampled (i.e., network-area relationships (NARs); (Dallas and Jordano, 2021)), which is a different aspect of the biodiversity of interactions (Galiana et al., 2021). The relationship between interaction richness and area sampled is expected to be positive, as larger areas are expected to include more species (Rosenzweig, M.L., 1995) and thus more interactions. However, this relationship is not linear, as it saturates at some point according to a species-area trend (Martinez, 1992; Galiana et al., 2021), and the number of interactions recorded in a given area is also dependent on the sampling effort (i.e., how many individuals are sampled in each species). It thus appears that the laws governing how interaction richness increases with partner richness (either species richness, in species-based networks, or number of distinct trees in a population, in individual-based networks) are analogous, with the latter being just a subset of the former. Unified scaling models, combining species richness and area sampled (Brose et al., 2004), are needed to fully understand the biodiversity of interactions in plant-frugivore assemblages. After all, scale-dependent co-occurrence factors of plants and frugivores govern the probabilities of their interspecific encounters, both at species and individuals scales and may yield accurate predictions of interaction occurrence across scales, from local populations and communities to metacommunities.

### Redundancy and functions in plant-frugivore interactions

The functional diversity of frugivore assemblages is expected to increase with increasing species richness, as more species are added to the assemblage (Cadotte et al., 2011). This is expected either for the frugivore assemblage of a plant species or the frugivore assemblage visiting a specific tree individual within a local population. However, such increase in functional diversity is not expected to be linear, as it saturates at some point (Schleuning et al., 2014; Donoso et al., 2017). In fact, the functional diversity of frugivore assemblages is often dominated by a few key species that provide most of the functional effects, while the rest of the species contribute little to the overall functional diversity (Rother et al., 2016). This pattern of functional redundancy is common in many ecological systems and has important implications for ecosystem functioning and resilience (Petchey and Gaston, 2002).

Here I examine the functional characteristics of frugivore assemblages using two species, *Prunus mahaleb* (Jordano and Schupp, 2000) and *Euterpe edulis* (Galetti et al., 2013), as case studies. I examine how functional diversity of frugivore assemblages increases with increasing species richness, and how this functional diversity is related to the functional traits of the frugivores. I use several functional diversity indices, including functional richness, functional evenness, functional divergence, and functional dispersion, to assess the functional diversity of the frugivore assemblages. First, I examine how variation in functional traits of frugivore species and the resulting interaction effectiveness generate gradients of interaction outcomes ranging from purely mutualistic to purely antagonistic effects on fitness. Second, I show how the functional diversity increases as the species richness of the frugivore assemblage increases. Data for functional traits come from (Jordano et al., 2025; Bovo et al., 2018; Galetti et al., 2013). My aim is not a comprehensive comparison of non-tropical and tropical assemblages, which would require including more case studies, but rather to illustrate how functional diversity increases with species richness and how this is related to the functional traits of the frugivores.

The ranked contributions of individual frugivore species in the two assemblages (Fig. 4) according to the percentage of mutualistic outcomes accounted for (i.e., frequency of fruit feeding instances including ingestion and/or carrying of seeds away from the plants) during their feeding visits shows a highly skewed pattern. There is a gradient ranging between extremes of species with 100% mutualistic interaction outcomes (no fruits dropped, or seeds damaged, during foraging) to those with purely antagonistic outcomes (100% of the feeding sequences ending in fruit dropping and/or seed predation). The former includes legitimate seed dispersers (Snow and Snow, 1988) with largely mutualistic outcomes in any interaction (e.g., *Sylvia* and *Curruca* warblers, *Turdus* spp. in *Prunus mahaleb*; large frugivores such as *Rhamphastos* spp., *Aburria jacutinga, Pyroderus scutatus, Procnias nudicollis*, and some *Turdus* in *Euterpe edulis*). The latter include largely granivorous species (e.g., psittacids) and fruit mashers (sensu Levey, 1987) such as thraupids and fringillids. Thus, both assemblages include three subsets of species: (1) species with purely mutualistic outcomes, (2) species with mixed outcomes (i.e., some feeding visits resulting in fruit dropping and/or seed predation), and (3) species with purely antagonistic outcomes. The latter are typically pulp consumers that consistently do not disperse seeds or do that very sporadically (e.g., fringillids), or seed predators that consume seeds without pulp.

Even among the most antagonistic species infrequent seed dispersal outcomes have been documented, in some cases resulting in non-trivial frequencies of actually mutualistic outcomes by species that otherwise would be categorized as pure antagonists (e.g., Blanco et al., 2016). Thus, even at the species level, frugivore assemblages are made up by a mixture of mutualistic and antagonistic interactions, with the relative contribution of each type of interaction varying among species in similar ways as shown previously for variation across species in, e.g., synzoochorous assemblages (Gómez et al., 2019). We can expect this heterogeneity in outcomes to be reflected in the functional diversity of the frugivore assemblages (Simmons et al., 2018). In fact, the functional diversity of frugivore assemblages of the two study species increases with increasing frugivore species richness (Fig. 3). The functional richness *FRic* and functional dispersion *FDis* both increase with increasing species richness, but the increase is not linear, as it saturates at some point (Fig. 3). This is consistent with previous findings that functional diversity saturates with increasing species richness (Schleuning et al., 2014; Donoso et al., 2017), a pattern shown in the trend of *FDis* in (Fig. 3) for the two frugivore assemblages.

**Figure 3:**
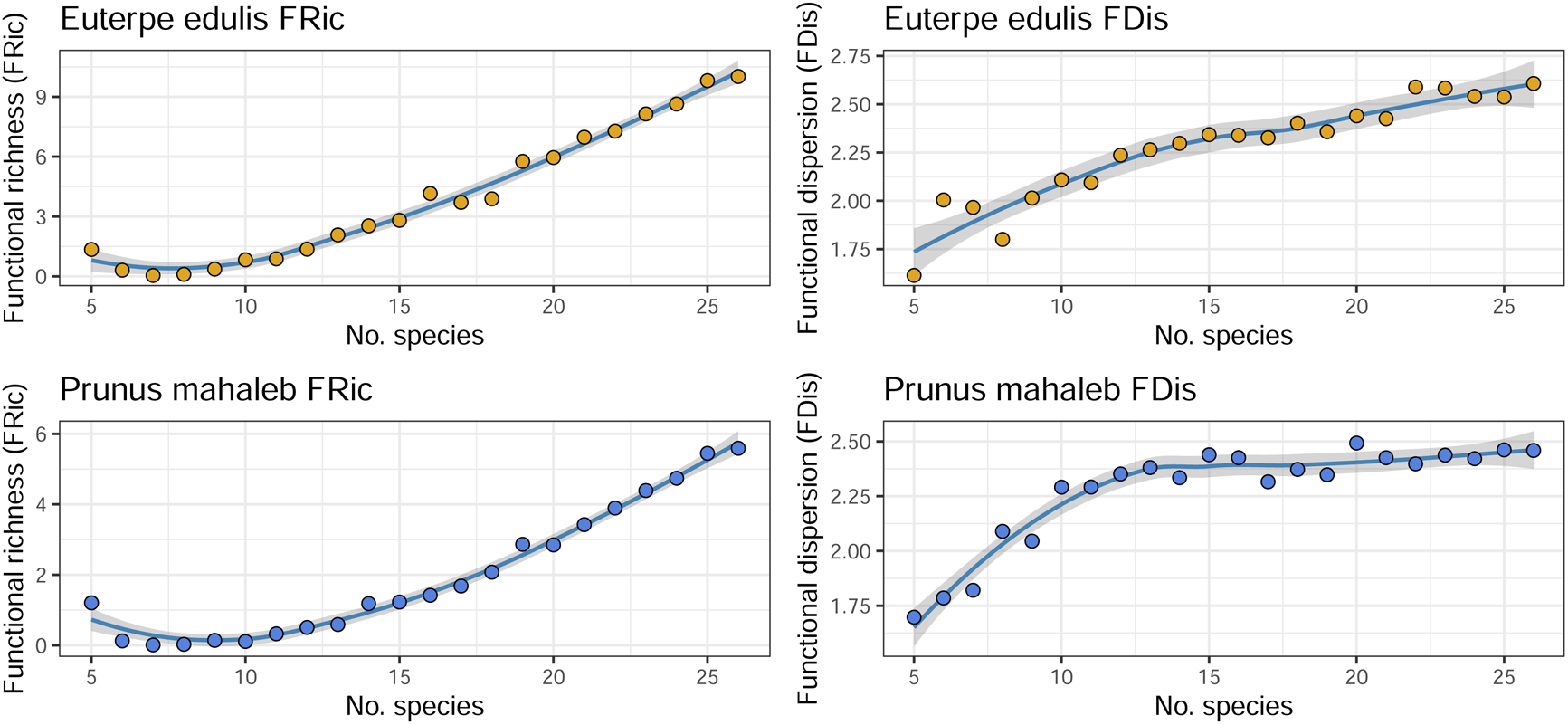
Variation of functional richness (*FRic*) and functional dispersion (*FDis*) of the *Prunus mahaleb* (blue) and *Euterpe edulis* (orange) frugivore assemblages as a function of cumulative number of species included in the assemblage. Intervals in grey around the smoothed splines indicate the range of values for the functional indexes when resampling the data (bootstrapping) to generate simulated assemblages with increasing number of species.

This is more evident for the non-tropical, *Prunus mahaleb* assemblage with *FDis* saturating in a range between *FDis* = 2.35 – 2.50 while for *Euterpe edulis FDis* steadily increases in the range *FDis* = 2.20 – 2.80 (Fig. 3). In terms of *FRic* the accumulation is steeper and reaches higher values for *Euterpe edulis*, as expected from its highly-diversified, tropical assemblage. In contrast, functional dispersion *FDis* does not show clear diverging trends between the two species with increasing species richness of the frugivore assemblages, suggesting a somehow similar range of functional roles. This is consistent with the idea that some species are more functionally redundant than others (Petchey and Gaston, 2002; Simmons et al., 2018), and that functional diversity is not just a simple function of species richness.

### Functional correlates of seed dispersal effectiveness

Considered from the above perspective, frugivore assemblages are not just a collection of species, but rather a complex system of interactions with distinct functional roles and outcomes. We may expect some covariation between functional roles (in terms of, e.g., seed dispersal effectiveness) and the contribution of each species to the functional diversity of the assemblage relate to its species-specific traits. In fact, the contribution of each species to the functional diversity of the frugivore assemblage is not evenly distributed, with some species contributing more than others (Fig. 4). In both assemblages species-specific seed dispersal effectiveness values (for the quantitative component of effectiveness, *SDE*_*Q*_) appear positively and significantly correlated with the contribution to functional diversity, *contrib*_*FD*_, of each species (*r* = 0.6047, *P* = 0.0001, *d*.*f*. = 47). Yet the correlation is not significant for *Euterpe edulis* (*r* =0.1353, *P* = 0.54, *d*.*f*. = 21) and highly significant for *Prunus mahaleb* (*r* = 0.8033, *P* ≪ 0.001, *d*.*f*. = 24) (Fig. 4)(also see Fig. Suppl. Mat. 1). These are two frugivore assemblages with highly diversified animal partners, and more complete comparative analyses of tropical and non-tropical assemblages are needed for proper generalization. Yet this result may suggest that the quantitative component of *SDE* maybe more responsive to variations in functional distinctness in non-tropical assemblages (i.e., more sensitive to variation in local abundance) than in the, even more diversified, tropical assemblages. In the latter, variation in the quantitative component of *SDE* is most likely more influenced by variation in fruit handling and processing than by variation in local abundance. Fruit/seed handling processing modes and behaviours are more diversified in tropical than in temperate species (Moermond and Denslow, 1985; Snow and Snow, 1988). It would be interesting to explore generalized trends for higher functional distinctness correlating with seed dispersal effectiveness with additional study species.

**Figure 4:**
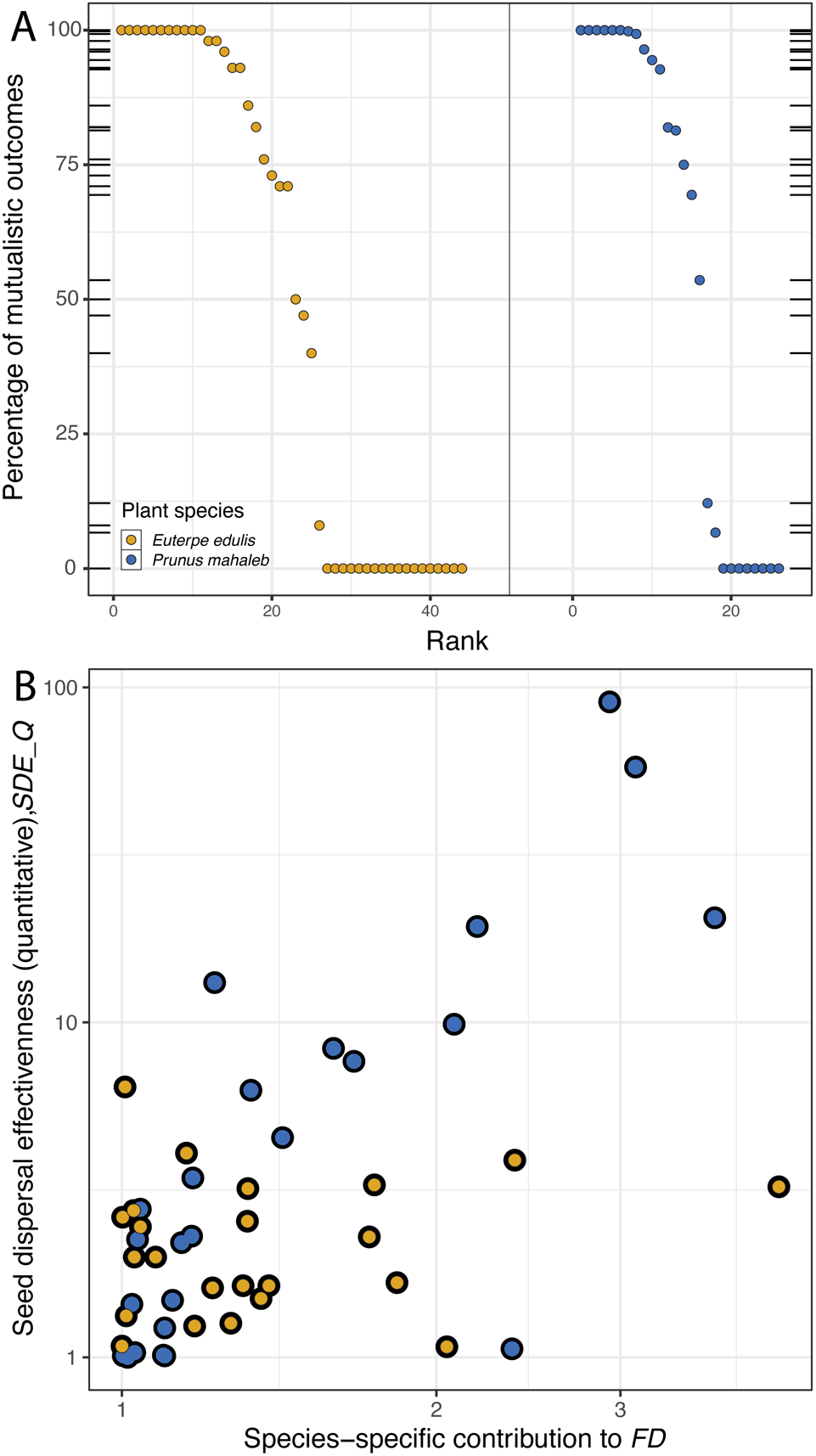
A. Distribution of plant-frugivore interaction outcomes among species in the frugivore assemblages of *Prunus mahaleb* (blue) and *Euterpe edulis* (orange). Points indicate individuals species, ranked according to the percentage of mutualistic outcomes (fruit feeding instances including ingestion and/or carrying of seeds away from the plants) during their feeding vists. Rugged tick marks along the Y axis indicate the position of species along this gradient between fully antagonistic (value= 0) to totally mutualistic (value= 100) outcome. B. Relationships between speciesspecific contbutions to functional diversity, *FD*, and the quantitive estimate of seed dispersal effectiveness (*SDE*) for *Prunus mahaleb* (orange) and *Euterpe edulis*.

In relation to species-specific traits determining variation in *FD*, the correlates among variables appear highly consistent for the two species assemblages (Fig. Suppl. Mat. 1, Table Suppl. Mat. 1). The most important traits covarying across species with any *SDE*_*Q*_ component (i.e., total effectiveness, per-visit effectiveness, and species contribution to *FD*) (Fig. Suppl. Mat. 1) include body mass (positively related to effectiveness per visit; *r* = 0.5585, *P* ≪ 0.0001, *d*.*f*. = 47), and the two proxies for local abundance (no. birds censused-birds.km- and visitation rate-vis.10h-Fig. S1), with the latter two being highly correlated (*r* = 0.8953, *P* ≪ 0.0001, *d*.*f*. = 47). Species-specific contributions to *FD*, also positively correlated with the two proxies for local abundance (no. birds censused-birds.km- and visitation rate-vis.10h-Fig. S1), appear significantly and positively associated to variation in total effectiveness and the resulting contribution to seed dispersal service (Fig. S1). These results suggest that species with higher contributions to assemblage *FD* show higher ^7^ contributions to *SDE*_*Q*_, either due to increased per-visit effectiveness or higher ^8^ visitation rates.

Previous research has shown that these generalized mutualisms between free-living frugivores and plants are largely based on resource-gathering interactions. Thus, in diversified assemblages of frugivores usually the most abundant species are those with higher contributions to *SDE* (Rother et al., 2016; Donoso et al., 2017) which shows highly skewed rankings of frugivore partners according primarily with local abundance (Quintero et al., 2025). This is consistent with the idea that frugivore assemblages are structured by the relative abundance of species, which in turn is influenced by the availability of resources (i.e., fruiting plants, (García et al., 2009)), determining interaction frequencies and the functional roles of each species in the assemblage. This is consistent with the idea that functional diversity is a complex trait that depends on both species richness and relative abundance (Cadotte et al., 2011). We may expect that in assemblages where variation in local abundance is not too large among species, the functional traits of the species might be more relevant in driving variation in *SDE*. The same can also be expected for frugivore assemblages in oceanic islands (Nogales et al., 2024), characterized by very high phylogenetic diversity and distinctness.

### Multiplexed interactions

A persisting challenge in the analysis of ecological networks has been the simultaneous consideration of multiple forms of interaction, such as the diverse range of antagonistic and mutualistic interactions occurring along the reproductive cycle of higher plants (Harper, 1977; Wang and Smith, 2002). All these myriad interactions-the plant’s interactome-are temporally overlaid throughout the reproductive cycle, previous to anthesis (e.g., bud formation) until well after seed dissemination and seedling establishment. This complex suite of interactions is difficult to disentangle, and recent research on complex networks has provided both conceptual and methodological advances, yet with limited application in mainstream ecological research (De Domenico et al., 2014; De Domenico, 2022). More generally, ecological systems can be represented as superpositions of different types of interaction networks involving multiple types and forms of interaction (Pilosof et al., 2017; Hutchinson et al., 2019), and deciphering the ecological processes behind this complexity is a lasting challenge.

I examine the role of mutualistic seed dispersal by animals within the multiplex structure of antagonistic and mutualistic interactions during the demographic cycle of a tree species, *Prunus mahaleb* (Rosaceae), built on individual-based interaction forms for different processes (e.g., herbivory, pollination, seed dispersal) (Fig. 1)(see the interaction networks involved in Suppl. Mat. Fig. 2).

The multiplex structure of the frugivore assemblage of *P. mahaleb* is illustrated in Fig. 5) where the interlayer links between the different layers may represent, e.g., the fitness consequences of each interaction type on the status of the plant nodes in successive layers. For example, the herbivory layer may affect the flowering and fruiting success of the plant nodes in the pollination layer, which in turn may affect the seed dispersal success in the seed dispersal layer. Such multiplexed structure is the result of overlaying multiple forms of interaction (Suppl. Mat. Fig. S2) which have been traditionally studied separately (Hutchinson et al., 2019). The repeated occurrence of the plant nodes in the different layers nicely portrays the sequential effects that multiple types of interaction have throughout every reproductive event. The multiplex structure allows us to visualize how the different interaction types are intertwined and how they contribute to the overall fitness of the plant species. Multiplexty (Bianconi, 2021) is thus a characteristic feature of individual-based interaction networks depicting the whole interactome of different plant individuals in a population (Fig. 5).

**Figure 5:**
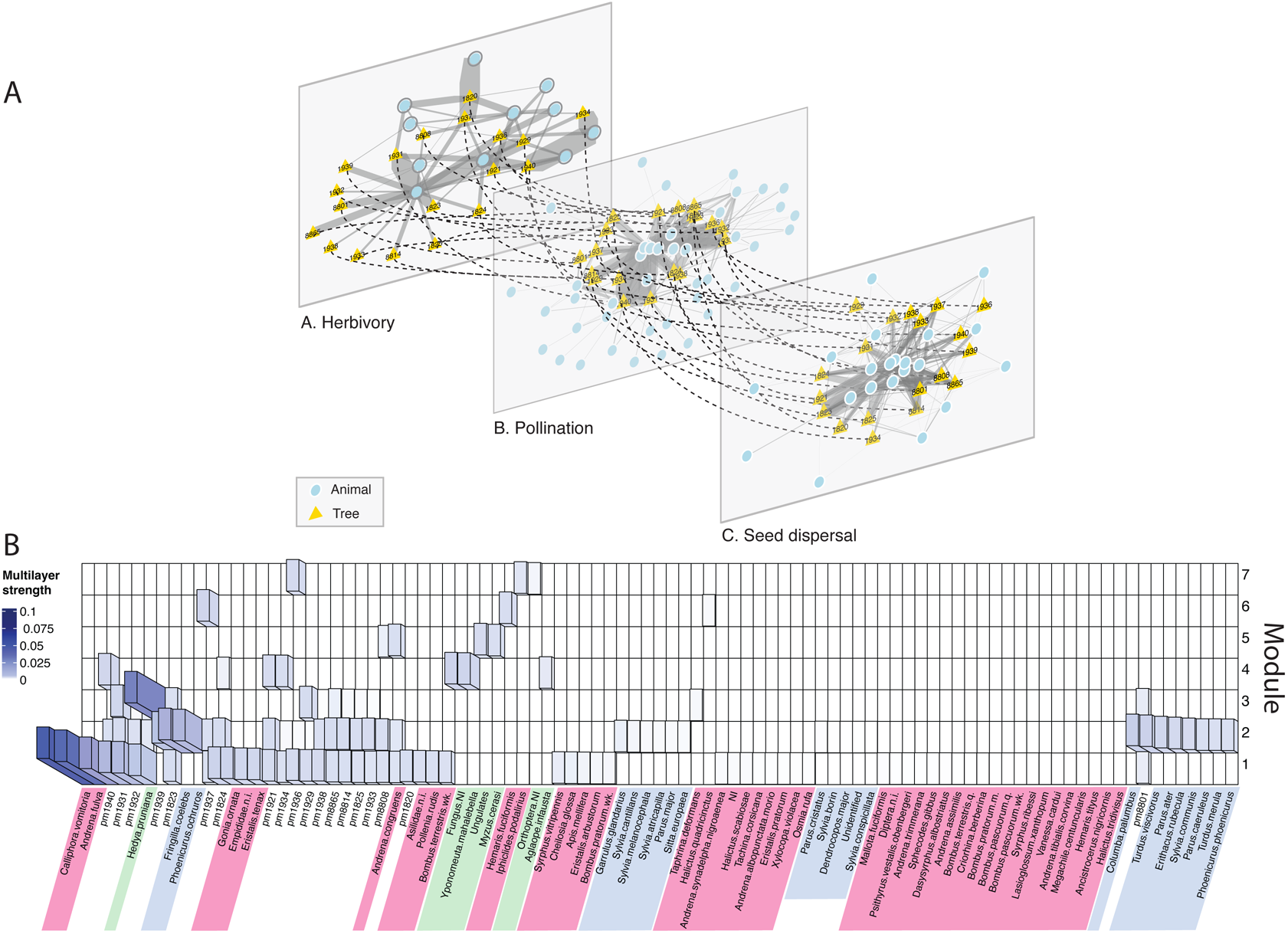
A. A multiplex network illustrating the interaction between invidiual *Prunus mahaleb* trees (yellow triangles) and their herbivore, pollinators, and seed disperser partners (blue circles). Dashed lines connect the same individual tree across layers, and represent the interlayer links illustrating the fitness consequences of each interaction type on the status of the plant nodes in successive layers. A Kamada-Kawai algorithm was used for the layout. B. The modular structure of the multiplex network illustrtaing 7 different modules and their composition including both trees and animal/fungi partners. Interaction strengths are standardized and illustrated as different blue shadings. Individual trees are identified by their field codes, with pm initials (e.g., pm1921, in white). Different colors group partners of different layers: herbivores (green), pollinators (red), and seed dispersers (blue).

**Figure 6:**
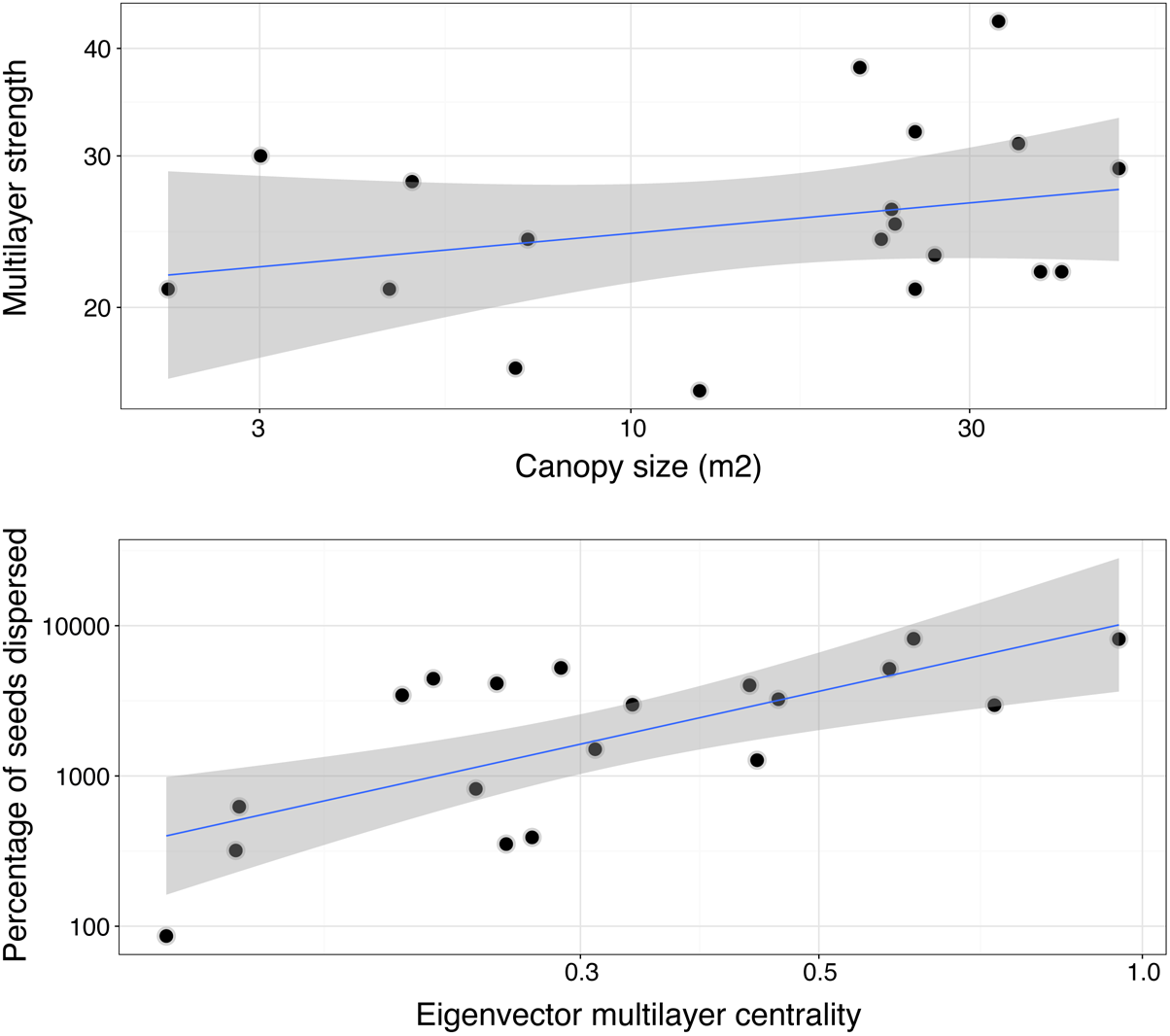
Correlates of multilayer strength (top) and fruit removal success (bottom, number of seeds succesfully dispersed from maternal tree) with canopy size and multilayer (eigenvector) centrality for *Prunus mahaleb* tress. Envelopes in grey indicate the 95% confidence intervals for the regression lines. The node strengths and centralities were estimated with Infomap (Edler et al., 2017; Frydman et al., 2023).

In our case study with *Prunus mahaleb*, I considered three types of interactions: (1) herbivory, including fungi and leaf and flower herbivores, (2) pollination, and (3) seed dispersal (Suppl. Mat. Fig. S2). Each of these interaction types is represented by a separate layer in the multiplex network, with the plant nodes appearing in each layer as state nodes (Fig. 5). Here I am not considering cumulative fitness effects but rather the sequential effects of each interaction type on the plant’s final fruit removal success, as the interlayer links are set to the same value. The multiplex structure allows us to visualize whether there is a significant signal of the plant-frugivore mutualism on top of the previous influences of all the biotic interactions occurring earlier in the reproductive cycle.

### Plant-frugivore interactions within the plant’s interactome

Multilayered networks have been extensively used to depict interactive sets of distinct networks, each embedding its own sets of interacting components (Kouvaris et al., 2015; De Domenico et al., 2016; Wang et al., 2016; De Domenico, 2022). Recent uses in ecology included their application to depict, e.g., multitrophic interactions in food webs, multilayered parasitism and host-parasitoid interactions, and integration of spatial and temporal slices of different types of ecological networks Kéfi et al. (2015); Pocock et al. (2016); Sauve et al. (2016); Pilosof et al. (2017); Hutchinson et al. (2019); Hervías-Parejo et al. (2024); Nogales et al. (2025). Multiplex networks are multilayer systems of *N* nodes that can be linked in multiple interacting and co-evolving layers (Menichetti et al., 2014). The multiplex structure of plant-animal interaction networks makes full biological sense when considering individual-based interaction networks (e.g., Rodríguez-Rodríguez et al., 2017) where individual plant nodes are repeated in distinct layers depicting the interactions with flower herbivores, pollinators, pre-dispersal seed predators, and seedling herbivores. Each plant node is connected across layers through the sequential fitness effects that emerge as outcomes of the different interaction types (Fig. 1).

The multiplex structure of the interactions involved in the three layers is represented in a supra-adjacency matrix (Suppl. Mat. Fig. 3) composed of three adjacency matrices as well as the inter-layer links. Two relevant aspects of the multiplex structure are worth noting: (1) the interlayer links are not weighted, as they represent the sequential fitness effects of each interaction type on the plant nodes in successive layers and are not empirically estimated in this study; yet, these were later estimated with Infomap (see Suppl. Mat.) for illustrative purposes. And (2) the plant nodes are repeated in each layer, while the animal/fungi taxa nodes are physical nodes appearing in each of the interaction types where the taxon participated. The multiplex structure thus allows a characterization of each tree’s centrality in the overall set of ecological interactions occurring in the study population. We might expect local trees with high eigenvector centrality to be more successful in terms of fitness, as they are more connected to the different interaction types and thus have higher chances of being visited by multiple partners. In fact, the reproductive success-in terms of seeds dispersed successfully-of each tree in the multiplex network is significantly correlated with its centrality (6, Suppl. Mat. Fig. 4). Obviously, the centrality of each tree in the multiplex network is not just a function of its interaction with frugivores, but also with other interaction types such as herbivory and pollination. This suggests that the overall fitness of a tree is determined by the cumulative effects of all the interactions it has with different partners, and that these interactions are interrelated in complex ways. Therefore, we might expect ample variation in the balance between mutualistic/antagonistic partners in different species and study systems (Hutchinson et al., 2019), with some systems being dominated by mutualistic interactions while others are dominated by antagonistic interactions.

The analysis also reveals a significant signal of tree size on the multilayer strength values of the trees in the multiplex network (*r* = 0.5992, *P* = 0.007, *df* = 17). In addition, larger trees are more likely to be visited by multiple partners and thus have higher chances of being successful in terms of seed dispersal (*r* = 0.6930, *P* = 0.001, *df* = 17, for the correlation between eigenvector centrality and number of seeds dispersed) (6, Suppl. Mat. Fig. 4). This is consistent with previous findings that larger trees have higher reproductive success (Jordano, 1995; García et al., 2009). Both the multilayer strength and centrality of each tree showed ample variation, which is more evident for strength (Suppl. Mat. Fig. 4). Only a few trees show high multilayer strength values (*pm*8801, *pm*1938, *pm*1929, *pm*1931), and its variation appears correlated mostly with strength in the seed dispersal interactions; yet, two of the trees with largest multilayer strength (*pm*1936, *pm*1929) had larger strength in the pollination interactions. These four trees are relatively large individuals, producing large crops of flowers and fruits, and growth in patches surrounded by other *P. mahaleb* trees. In general, the interaction strength with herbivores is lower across all the trees when compared with strength values for pollination and, especially, seed dispersal. Yet those trees with lowest multilayer strength (*pm*1921, *pm*1937) show the largest strength with herbivores. The position and global interaction strength of each tree in the multiplex network is thus a function of its size, its interaction with frugivores, and its interaction with other partners such as herbivores and pollinators.

In relation with centrality (Suppl. Mat. Fig. 4), only three trees (*pm*1940, *pm*1931, *pm*1934) had largest eigenvector centrality yet none of them (except *pm*1931) showed a large multilayer strength, suggesting that multilayer centrality and strength appear decoupled when considering the whole population. The reason for this decoupling most likely relates to how the strength and centrality values of each tree in the monolayer networks covary or not. And it appears they don’t (Suppl. Mat. Fig. 5), at least for eigenvector centrality. In fact, eleven trees with largest eigenvector centrality (*EC*) values in the multiplex network (*pm*1940, *pm*1931, *pm*1934, *pm*1823, *pm*1921, *pm*1936, *pm*1937, *pm*1929, *pm*19281, *pm*8801, *pm*1823) (above diagonal in Suppl. Mat. Fig. 5) show reduced centrality values in the monolayer networks (Suppl. Mat. Fig. 5). This suggests that the centrality of each tree in the multiplex network is not just a function of its interaction with frugivores, but also with other interaction types such as herbivory and pollination. Trees with a combination of high centrality values in two or more monolayer networks (e.g., *pm*1940, *pm*1931, *pm*1823) show high multilayer centrality values, suggesting that the overall centrality of a tree in the population interactome is determined by the cumulative effects of all the interactions it has with different partners, and that these interactions are interrelated in complex ways.

The pattern for multilayer strength, in contrast, appears with a greater covariation with the strength in the monolayer networks (Suppl. Mat. Fig. 5). In fact, the strength of each tree in the multiplex network is significantly correlated with its strength in the seed dispersal layer (*r* = 0.707, *P* ≪ 0.0001, *df* = 47), and also with its strength in the pollination layer (*r* = 0.603, *P* ≪ 0.0001, *df* = 47), but not with its strength in the herbivory layer (*r* = 0.135, *P* = 0.34, *df* = 47). This suggests that the strength of each tree in the multiplex network is primarily determined by its interaction with a specific set of partners, especially frugivores and/or pollinators, while its interaction with herbivores has a lesser effect on its overall strength.

The modularity analysis, based on the Infomap algorithm for multilayer networks (Edler et al., 2017; De Domenico, 2022), was carried out on the supra-adjacency matrix (see Suppl. Mat.) and reveals the presence of seven distinct modules in the multiplex network 5. Each module is composed of a subset of plant nodes and their associated animal/fungi partners across the three layers. The modules are not evenly distributed across the layers, with some modules being more prominent in certain layers than others. For example, module 1 is primarily composed of plant nodes and their associated partners in the pollination layer, while module 2 is primarily composed of plant nodes and their associated partners in the seed dispersal layer. Modules 1-2 include the animal/fungi taxa with higher interaction frequencies in the tree population. The rest of modules, 4-7 appear closely associated with trees with higher frequency of interactions with herbivores/fungi (e.g., *pm*1937, *pm*1940, *pm*1936, *pm*8808). This suggests that different interaction types may have distinct modular structures, which may reflect the different ecological processes underlying each interaction type.

Tree size, and other tree traits related to size effects such crop size of flowers and fruits, appear as major drivers of the position of each tree in the population interactome, with a significant signal on realized fitness in terms of successfully-dispersed seeds 6. The strength and centrality of each tree in the multiplex network is thus a function of its size and size-related fecundity characteristics that may determine variable interaction frequencies with frugivores, as well as with other partners such as herbivores and pollinators. This suggests that the overall fitness of a tree is determined by the cumulative effects of all the interactions it has with different partners, and that these interactions are interrelated in complex ways.

The central role of size and hierarchies in plant populations have been extensively documented, especially for trees (Harper, 1977; Weiner and Solbrig, 1984; Weiner, 1988). It would be interesting to consider how this role of tree size in structuring an individual’s tree interactome changes along the ontogeny of the tree, especially for long-lived tree species (Petit and Hampe, 2006). We might expect extensive variation along trees ontogenies in the accumulated diversity of interactions and their lasting effects, so that some individuals get increased benefits from diversified interactomes and accumulate gains along their lives resulting in higher centralities and strength in the population. It remains to be tested how such ontogenetic trajectories of individual interactomes mirror the patterns highlighted in this transversal study, i.e., to obtain a description of the ontogeny of an individual’s interactome.

Our study with *Prunus mahaleb* demonstrated an extended effect of size hierarchies on the structure of local population interactomes, with consequences for plant fitness and points for important implications in plant demography and applied aspects of forest restoration. For example, conservation of large trees is fundamental for the prospects of local population growth and recruitment, as they have the largest contributions to *λ* (Condit et al., 1998). Large trees, because of their significant centrality and strength in the multiplex structure of the interactions involved in each reproductive cycle (at least in the three layers of herbivory, pollination, and seed dispersal), should be key targets in preservation actions (Lindenmayer et al., 2012), as their significant demographic effects pivot on their distinct role in sustaining the biotic interactomes that guarantee forest regeneration.

## Concluding remarks

The size and diversity of plant-frugivore interactomes (i.e., the whole suite of interactions involved in the seed dispersal system of higher plants) steeply increases with partner diversity, in parallel to a marked increase in the functional diversity of frugivore assemblages. Such functional diversity is not just a function of species richness, but also of the relative abundance of species and their functional traits. This suggests that frugivore assemblages are primarily structured by the relative abundance of species, which in turn is determined by the availability of resources (i.e., fruiting plants). This is expected from the generalized, resource-consumer, interactions among the free-living partner species involved in these complex networks. A central issue in the analysis of interaction biodiversity is thus to determine which elements (species, interactions) of these highly diversified interactomes are more relevant in contributions to the overall functional outcomes of these mutualistic assemblages. When considered in combination with the myriads of other biotic interactions occurring during each reproductive cycle of the higher plants, animal-mediated dispersal has a central role as a driver of variation in seed dispersal success and realized fecundity.

Our study on *Prunus mahaleb* suggests that size hierarchies exert a prolonged influence on the structural organization of local population-scale interactomes, with consequent effects on plant fitness and notable implications for both population demography and the applied domains of forest restoration. In particular, the conservation of large individual trees emerges as a critical factor for sustaining local population growth and recruitment, given their disproportionately high contributions to the population growth rate. Owing to their pronounced centrality and interaction strength within the multiplex network encompassing key reproductive processes-specifically herbivory, pollination, and seed dispersal-large trees should be considered priority targets in conservation and management strategies.

## Acknowledgements

I appreciate the help in the field and extensive discussions and ideas of Eugene W. Schupp, Cristina García, Myriam Márquez, JuanLu García-Castaño, Alfredo Valido, Jesú s G.P. Rodríguez, Jordi Bascompte, Paco Rodríguez-Sánchez, Blanca Arroyo-Correa, Miguel Jácome-Flores, JuanPe González-Varo, Elena Quintero, Jorge Isla, Irene Mendoza, Gemma Calvo, Juan Mi Arroyo, and many others. I’d like to highlight the sustained help, enthusiasm and generosity of my wife, Myriam Márquez, and the late Manolo Carrión and JuanLu García-Castaño. Field work with *Prunus mahaleb* was greatly assisted by Manolo Carrión, Eugene W, Schupp, JuanLu García-Castaño, Cristina García, and especially Myriam Márquez. Over the years my field work was generously supported by the facilities at Parque Natural de las Sierras de Cazorla, Segura y Las Villas (Junta de Andalucía), ICTS-Reserva Biológica de Doñana (CSIC) and the administration of the Doñana National Park. Completion of this manuscript was funded by grant PID2022-136812NB-I00 from the Agencia Estatal de Investigación, the Plan Propio de Investigación y Transferencia, University of Sevilla (2021-2024), and a LifeWatch ERIC-SUMHAL project (LIFEWATCH-2019-09-CSIC-13) with FEDER-EU530 funding.

