## Supplementary Material for "The biodiversity of plant-frugivore interactions: types, functions, and consequences"

OIKOS
ADVANCING ECOLOGY

### Supplementary Material

### The biodiversity of plant-frugivore interactions: types, functions, and consequences

Pedro Jordano[1,2]

Sevilla, November 11, 2025

[1] Integrative Ecology Group, Estación Biológica de Doñana, CSIC, Avda. Americo Vespucio 26, E-41092 Sevilla, Spain.

[2] Dept. Biología Vegetal y Ecología, Universidad de Sevilla, Sevilla, Spain.
Suppl. Material: datasets (both raw and summarized) and code used in any analyses.
Support repository in GitHub.
(https://github.com/pedroj/MS_Oikos_SI-FSD2024)

### *Prunus mahaleb* and *Euterpe edulis* data

*Prunus mahaleb* (Rosaceae) is a shrub or small tree native to the Mediterranean region. It is a keystone species in the Mediterranean forest, where it is dependent on birds for seed dispersal. The species has been introduced to North America and South America, where it is cultivated for its edible fruits and seeds. Data for the species (frugivory and seed dispersal) come from Jordano (1995); Jordano and Schupp (2000). Information on pollinators was obtained from (Jordano, 1993). The tree is gynodioecious, with both female and hermaphroditic individual plants. Flowers are visited by a diversified assemblage of bees and flies totaling more than 40 species. Flies are more frequent visitors to male-sterile trees than bees. Herbivory was assessed for each tree by visual censuses and counts of damaged branches and leaves. For each tree, 3-5 different areas (0.5 x 0.5 m) of the canopy at different heights were scanned and the number of branches with ungulate marks was counted. In addition, leaves were inspected for fungi presence, with visual identification, when possible, of the taxa involved based on damage type, form, coloration, etc. A relative damage score was assigned (0-5 ranging from absence through sporadic, frequent, common, and extreme damage) based on counts. Herbivorous insects were censused visually at each tree during the herbivory scans and a relative damage score was also assigned (0-5 ranging from absence through sporadic, frequent, common, and extreme damage) based on counts. Herbivory surveys were carried out from the initial leafing stages until leaves were expanded at full size. This period encompassed the flowering period and extended 2-3 weeks after flowering. Regarding the seed dispersal stage, the tree is dispersed by 38 bird species, including thrushes (*Turdus* spp.), robin and redstarts (*Phoenicurus ochruros, Erithacus rubecula*), and several species of finches (*Fringilla* spp.) and parids

*Parus, Cyanistes* spp. feed on pulp and sporadically disperse seeds. Among them seed disperser species account for 42.2% of the records, pulp consumer species for 17.2%, and pulp consumer species that sporadically disperse some seeds, for 9.4% of all the species recorded.

The palmito or juçara palm (*E. edulis*, Arecaceae) is a dominant palm species endemic to the Atlantic forest and dependent mostly on birds for successful seed dispersal. It also occupies fragmented forest stands, remnants within the past 200 years since the establishment of extensive coffee plantations in São Paulo state (Galetti et. al. , 2013). Data for *E. edulis* come from (Galetti et. al. , 2013). Fifty-eight birds and 21 mammal species have been recorded feeding on the fruits of *E. edulis* in its distribution range. Of them, only 32 bird and six mammal species can act as legitimate seed dispersers (animals that regurgitate or defecate the seeds). The main seed dispersers are birds ranging from small thrushes (*Turdus* spp.) to large toucans (*Ramphastos* spp.). Smaller thraupids and tyrannids (e.g., *Euphonia* spp., *Tangara* spp.) feed on fruits but always drop them beneath the parent palm with no seed ingestion. Several parrot and parakeet species (e.g., *Pionus maximiliani*, *Triclaria malachitacea* and *Pyrrhura frontalis*), mid-size rodents (e.g., *Cuniculus paca* and *Dasyprocta* spp.), several species of small rodents, deer and peccaries are seed predators, and six mammals (marsupials, larger meso-mammals, tapir, etc.) are infrequent consumers of the fruits and may contribute infrequent seed dispersal events.

#### Sampling interactions

Frugivory events in *P. mahaleb* and *Euterpe edulis* were sampled in the field using the same protocol. We recorded all interactions between plants and frugivores (fruit consumers) for sampled trees in areas covering between 25 and 300 ha, respectively. The sampling was conducted during the fruiting seasons of *P. mahaleb* (late July to early September) and *E. edulis* (late January to mid June). We used a combination of direct observations at focal trees and feeding records during walked transects. For each interaction, we recorded the species involved, the type of interaction (active feeding and ingestion of fruit and seed, pecking pulp, crack-opening he seed, etc.), and the number of individuals involved (for details see Jordano and Schupp , 2000; Galetti et. al. , 2013). Sampling of direct watches at *P. mahaleb* trees totaled 190.3 and 165.0 tree-hours in two successive years (1988-1989) with additional sampling for ca. 60.0 tree-hours in 1990-1992 in four different populations. Feeding observations of frugivores at *E. edulis* were carried out during 2326 h of direct watches in ten different sites and 5-20 individual palms per site.

#### Combining diversified interactions into multiplex networks

We used a multiplex network approach (De Domenico , 2022) to combine the different types of interactions into a single network. The multiplex network is a type of network that consists of multiple layers, where each layer represents a different type of interaction. The multiplex network was constructed by creating a bipartite graph for each type of interaction, where the nodes in one set represent the species involved in that interaction type (partners), and the individual trees in the population as another set; the edges represent the interactions between them (Fig. S1). Thus, a multiplex network is a specific type of multilayer network where the same set (or subset) of nodes (the trees in our case) appears in every layer, but the edges in each layer represent different types of relationships (e.g., herbivory, pollination, and seed dispersal partners in each of three layers) that taken together characterize the interactome of the trees (Jordano, P. , 2024). Thus, multiplexed networks are ideal representations for the multiple, distinct types of interactions involved in every reproductive episode of the tree population.

In multiplex networks, interlayer edges only connect corresponding nodes across layers (i.e., a tree node in one layer is connected to its counterpart representation in another layer). There are no interlayer edges between different nodes. The nodes represent the same entities across all layers. For example, a tree could be represented in multiple layers corresponding to different biotic interaction types (e.g., with its herbivory, pollination, and seed dispersal animal/fungi partners). These partner nodes in turn are just present in the layer corresponding to their interaction type. Therefore, interlayer links among repeated instances of a given tree in the three layers may indicate cumulative fitness effects derived from a layer and with effects on the subsequent layer. For example, outcomes of interactions in the pollination layer have lasting consequences for the seed dispersal layer in terms of fruit set success.

Technically, a multilayer network is a quadruple $M = (AM, EM, V, L)$ (De Domenico et. al. , 2014; Kivelä et. al. , 2014; Boccaletti et. al. , 2014; De Domenico , 2022), where $AM$, $EM$ and $L$ are the nodes, links (interactions) and layers respectively, of the $M$ network, so that $(V, EM)$ is a graph and $V \subset AxL$. Multilayer networks are hypergraphs, which contain multiple graphs (in our case, networks) linked to each other by interlayer links. The tensor notation is currently used to represent such multilayer structures: $\tau_{j,\beta}^{i,\alpha} = M_{j,\beta}^{i,\alpha}, i, j \in 1, 2, ..., N$ and $\alpha, \beta \in 1, 2, 3$ for $L = 1, 2, ..., M$ layers. Multilayer networks can have several "aspects" in each layer (De Domenico , 2022), and an "elementary layer" is a single element in an aspect of stratification. For example, in Fig. 6, each layer is a "still photo" of interactions that actually occur over variable periods of time. Pollination interactions occur throughout the phenophase and we could imagine different variants of the $L_2$ layer corresponding to different months during flowering, and consider studies of the temporal dynamics of interaction networks (Olesen, JM et. al. , 2008)(see also Carnicer et. al. , 2009). Thus, each aspect of the network could contain the interactions observed in a certain time interval. Such relations are included using sequences $L = L_a d_a = 1$ of $L_a$ sets of elementary layers, where $a$ indexes the different aspects, $d$. We must take into account that $d = 0$ for a single-layer network; $d = 1$ when there is a single type of layer; and $d = 2$ when there are two types of layers. There may be different types of multilayer networks depending on whether there are links between layers and how they are arranged (see De Domenico , 2022): multiplexed or multiplexed, colored, general, etc. These configurations are very useful in ecology, where it is possible to represent variations of interaction networks in time or space (Pilosof et. al. , 2017; Hutchinson et. al. , 2019).

The current mathematical notation that allows greater precision in the analysis of multilayer networks can be found in Bollobás (1998), and De Domenico et. al. (2014); De Domenico (2022). It is based on the use of successive order tensors, which allow indexing the complex multidimensional structure of multilayer networks. Scalars ($a$), vectors ($a_i$) and matrices ($A_{ij}$) can be recognized by their simple tensors of rank 0, 1, and 2 respectively. Thus, an adjacency matrix for a classic network, single-layer, would have a tensor of rank 2 (two dimensions), $\tau_{ij\alpha} = a_{ij}^{[\alpha]}$, while a multilayer would have a multilayer tensor of adjacency of rank 3, $\tau_{i\beta}^{i\alpha} = M_{i\beta}^{i\alpha}, i, j \in 1, 2, ..., N$ and $\alpha, \beta \in 1, 2, 3$ for $L = 3$ layers, and a multilayer with aspects, a tensor of rank 4 (e.g., Jordano, P. , 2024).

Fig. 6 considers a special type of multilayer network (a multiplex) where plant nodes are present recurrently in all layers; these in turn represent interactions with herbivores (in vegetative parts), pollinators, frugivores and post-dispersive seed predation, all of them occurring sequentially. The multiplex network, G, obviously contains multiple networks in several layers, M, with N nodes ($G = G^{[1]}, G^{[2]}, G^{[3]}, ..., G^{[M]}$), so that each layer $\alpha = 1, 2, 3...M$ contains a network $G = V, E\alpha$, with its nodes $V$ and its set of $E\alpha$ bonds. Therefore, we will have different networks (plants by animals) for each type of interaction in each layer (Fig. S2), and networks with different types of interaction will be connected through the shared plants. That is, a representation $A = a_{[1]}, a_{[2]}, a_{[3]}, ..., a_{[M]}$ of different adjacency matrices that contain the interactions in each layer, $a_\alpha$ (Bianconi , 2021). The layers are "diagonally coupled", so the links between layers occur only between shared plants (Figs. 1, 5, S3), which represents the successive effects of the mutualists and antagonists on the individual fitness of the plants. The "aspects" referred to above would represent, for example, different networks corresponding to different time intervals, such as the seasons, of each of the types of interaction: this is the case for four aspects for L1 corresponding to herbivory interactions in each of the four seasons of the year.

In Fig. 6 only one aspect has been represented for each type of interaction, which is illustrated with a network in each layer. The analysis of multilayer networks exploits, for example, the interlayer degree correlations that exist between the degrees (degree, $k$) of each node in the different layers (De Domenico et. al. , 2014; Bianconi , 2021). The multiplex degree is not a scalar (as in a single-layer network), but is a vector: $(k_i = k^{[1]}, k^{[2]}, ..., k^{[M]})$, whose $k^\alpha$ elements are the degree values in each layer. But in multiplex networks the nodes can also be connected through the layers (Fig. 6) by means of the interlayer links. For example, in Fig. 6 the individual plants are connected through links that represent the fitness effects derived from the interactions in each layer. These links between layers are called multilinks: $\overrightarrow{m} = m^{[1]}], m^{[2]}, ..., m^{[M]}$, where the elements $m^\alpha \in 0, 1$ and are initially set to $m_\alpha = 2.0$ and later estimated with Infomap (Edler et. al. , 2017; Frydman et. al. , 2023). Therefore, multilayer networks are characterized by containing four elements (and not just two, like single-layer networks): nodes, links, multilinks and layers. In addition, nodes are divided into two groups, those that only connect with other nodes of their layer and those that have multilinks with nodes of other layers (for example, note that only individual tree nodes in Fig. S5A have multilinks). Thus in Fig. 6 only the tree nodes have multilinks. There are no multilayer solutions yet available for the analysis of bipartite networks, with which most of the analyses carried out so far refer to projections of the supra-adjacency matrix that represents all interactions in all layers, that is, in the form of matrices of added adjacency (see Frydman et. al. , 2023).

Table S1. List of datasets used in this study. Code, individual code for reference; Type, whether it is an individual-based (ind) or species-based network; $P$, number of plant species; $A$, number of animal species; $S$, total number of species; $I$, number of distinct pairwise interactions, $C$, connectance ($C = I/(A * P)$); Biome, biome type, t= tropical; nt= non-tropical; Site, specific study site. For species-based networks, species name is set to NA.

| ID | Code | Type | Species | $P$ | $A$ | $S$ | $I$ | $C$ | Biome | Reference |
|---|---|---|---|---|---|---|---|---|---|---|
| 1 | indw01 | ind | *Pistacia lentiscus* (Anacardiaceae) | 40 | 27 | 67 | 392 | 0.3630 | nt | Quintero et. al. 2023. |
| 2 | indw02 | ind | *Pistacia lentiscus* (Anacardiaceae) | 40 | 16 | 56 | 134 | 0.2094 | nt | Quintero et. al. 2023. |
| 3 | indw03 | ind | *Juniperus phoenicea* (Cupressaceae) | 35 | 10 | 45 | 137 | 0.3914 | nt | . Isla et. al. 2023. |
| 4 | indw04 | ind | *Juniperus phoenicea* (Cupressaceae) | 35 | 10 | 45 | 154 | 0.4400 | nt | Isla et. al. 2023. |
| 5 | indw05 | ind | *Juniperus phoenicea* (Cupressaceae) | 35 | 11 | 46 | 148 | 0.3844 | nt | Isla et. al. 2023. |
| 6 | indw06 | ind | *Lithraea molleoides* (Anacardiaceae) | 13 | 10 | 23 | 37 | 0.2846 | t | Vergara-Tabares et. al. 2022. |
| 7 | indw07 | ind | *Lithraea molleoides* (Anacardiaceae) | 14 | 10 | 24 | 33 | 0.2357 | t | Vergara-Tabares et. al. 2022. |
| 8 | indw08 | ind | *Lithraea molleoides* (Anacardiaceae) | 14 | 13 | 27 | 46 | 0.2527 | t | Vergara-Tabares et. al. 2022. |
| 9 | indw09 | ind | *Lithraea molleoides* (Anacardiaceae) | 13 | 12 | 25 | 41 | 0,2628 | t | Vergara-Tabares et. al. 2022. |
| 10 | indw10 | ind | *Lithraea molleoides* (Anacardiaceae) | 12 | 11 | 23 | 29 | 0.2197 | t | Vergara-Tabares et. al. 2022. |
| 11 | indw11 | ind | *Lithraea molleoides* (Anacardiaceae) | 11 | 7 | 18 | 25 | 0.3247 | t | Vergara-Tabares et. al. 2022. |
| 12 | indw12 | ind | *Laurus nobilis* (Lauraceae) | 18 | 17 | 35 | 87 | 0.2843 | nt | Rodríguez-Sánchez, F. 2010. |
| 13 | indw13 | ind | *Prunus mahaleb* (Rosaceae) | 19 | 20 | 39 | 211 | 0.5553 | nt | Jordano and Schupp 2000. |
| 14 | indw14 | ind | *Euterpe edulis* (Arecaceae) | 17 | 9 | 26 | 31 | 0.2026 | t | Friedemann et. al. 2022. |

| ID | Code | Type | Species | $P$ | $A$ | $S$ | $I$ | $C$ | Biome | Reference |
|---|---|---|---|---|---|---|---|---|---|---|
| 15 | indw15 | ind | *Euterpe edulis* (Arecaceae) | 15 | 7 | 22 | 25 | 0.2381 | t | Friedemann et. al. 2022 |
| 16 | indw16 | ind | *Euterpe edulis* (Arecaceae) | 30 | 8 | 38 | 50 | 0.2083 | t | Friedemann et. al. 2022 |
| 17 | indw17 | ind | *Cecropia glaziovii* (Urticaceae) | 27 | 37 | 64 | 124 | 0.1241 | t | Jordano, P. 2024a |
| 18 | indw18 | ind | *Heynea trijuga* (Meliaceae) | 24 | 11 | 35 | 48 | 0.1818 | t | Gopal et. al. 2020 |
| 19 | indw19 | ind | *Myristica dactyloides* (Myristicaceae) | 25 | 7 | 32 | 45 | 0.2571 | t | Gopal et. al. 2020 |
| 20 | indw20 | ind | *Persea macrantha* (Lauraceae) | 32 | 21 | 53 | 186 | 0.2768 | t | Gopal et. al. 2020 |
| 21 | indw21 | ind | *Henriettea succosa* (Melastomataceae) | 18 | 22 | 40 | 77 | 0.1944 | t | Crestani et. al. 2019 |
| 22 | indw22 | ind | *Prestoea decurrens* (Arecaceae) | 31 | 9 | 40 | 100 | 0.3584 | t | Lamperty et. al. 2021 |
| 23 | indw23 | ind | *Corema album* (Ericaceae) | 24 | 15 | 39 | 129 | 0.3583 | nt | Villalva et. al. 2024 |
| 24 | indw24 | ind | *Bursera penicillata* (Burseraceae) | 15 | 11 | 26 | 43 | 0.2606 | t | Ramaswami et. al. 2017 |
| 25 | indw25 | ind | *Erythroxylum monogynum* (Erythroxylaceae) | 12 | 6 | 18 | 29 | 0.4028 | t | Ramaswami et. al. 2017 |
| 26 | indw26 | ind | *Flacourtia indica* (Salicaceae) | 13 | 5 | 18 | 33 | 0.5077 | t | Ramaswami et. al. 2017 |
| 27 | indw27 | ind | *Flueggea leucopyrus* (Phyllanthaceae) | 10 | 8 | 18 | 22 | 0.2750 | t | Ramaswami et. al. 2017 |
| 28 | indw28 | ind | *Canthium coromandelicum* (Rubiaceae) | 10 | 8 | 18 | 30 | 0.3750 | t | Ramaswami et. al. 2017 |
| 29 | indw29 | ind | *Santalum album* (Santalaceae) | 14 | 10 | 24 | 38 | 0.2714 | nt | Ramaswami et. al. 2017 |
| 30 | indw30 | ind | *Ziziphus oenopolia* (Rhamnaceae) | 15 | 13 | 28 | 102 | 0.5231 | nt | Ramaswami et. al. 2017 |
| 31 | indw31 | ind | *Chamaerops humilis* (Arecaceae) | 39 | 6 | 45 | 76 | 0.3248 | nt | Jácome-Flores et. al. 2020 |
| 32 | indw32 | ind | *Chamaerops humilis* (Arecaceae) | 24 | 6 | 30 | 57 | 0.3958 | nt | Jácome-Flores et. al. 2020 |

| ID | Code | Type | Species | $P$ | $A$ | $S$ | $I$ | $C$ | Biome | Reference |
|---|---|---|---|---|---|---|---|---|---|---|
| 33 | indw33 | ind | *Miconia irwinii* (Melastomataceae) | 15 | 9 | 24 | 59 | 0.4370 | t | Guerra et. al. 2017. |
| 34 | indw34 | ind | *Juniperus macrocarpa* (Cupressaceae) | 26 | 11 | 37 | 72 | 0.2517 | nt | Villalva et. al. 2024. |
| 35 | indw35 | ind | *Prosopis flexuosa* (Fabaceae) | 26 | 9 | 35 | 72 | 0.3077 | nt | Miguel et. al. 2018. |
| 36 | indw36 | ind | *Prosopis flexuosa* (Fabaceae) | 28 | 10 | 38 | 84 | 0.3000 | nt | Miguel et. al. 2018. |
| 37 | indw37 | ind | *Prosopis flexuosa* (Fabaceae) | 54 | 10 | 64 | 112 | 0.2074 | nt | Miguel et. al. 2018. |
| 38 | indw38 | ind | *Prosopis flexuosa* (Fabaceae) | 35 | 8 | 43 | 61 | 0.2179 | nt | Miguel et. al. 2018. |
| 39 | indw39 | ind | *Prosopis flexuosa* (Fabaceae) | 29 | 9 | 38 | 99 | 0.3793 | nt | Miguel et. al. 2018. |
| 40 | indw40 | ind | *Schinus terebinthifolia* (Anacardiaceae) | 26 | 16 | 42 | 93 | 0.2236 | t | Vissoto et. al. 2022. |
| 41 | indw41 | ind | *Phillyrea angustifolia* (Oleaceae) | 10 | 16 | 26 | 72 | 0.4500 | nt | Moracho, E et. al. 2025. |
| 42 | indw42 | ind | *Phillyrea angustifolia* (Oleaceae) | 9 | 12 | 21 | 41 | 0.3796 | nt | Moracho, E et. al. 2025. |
| 43 | indw43 | ind | *Marcgravia longifolia* (Marcgraviaceae) | 24 | 43 | 67 | 127 | 0.1231 | t | Thiel et. al. 2024. |
| 44 | indw44 | ind | *Osyris lanceolata* (Santalaceae) | 19 | 14 | 33 | 62 | 0.2331 | nt | Moracho, E et. al. 2025. |
| 45 | indw45 | ind | *Naringi crenulata* (Rutaceae) | 22 | 12 | 34 | 62 | 0.2348 | t | Jayanth et. al. 2024. |
| 46 | indw46 | ind | *Ziziphus oenopolia* (Rhamnaceae) | 20 | 13 | 33 | 57 | 0.2192 | nt | Jayanth et. al. 2024. |
| 47 | sdw00 | sp | NA | 77 | 68 | 145 | 203 | 0.0388 | nt | Jordano, P. 2007. |
| 48 | sdw01 | sp | NA | 25 | 36 | 61 | 253 | 0.2811 | nt | Moracho, E et. al. 2025. |
| 54 | sdw02 | sp | NA | 29 | 78 | 107 | 479 | 0.2118 | nt | Moracho, E et. al. 2025. |
| 50 | sdw02a | sp | NA | 16 | 44 | 60 | 201 | 0.2855 | nt | Quintero et. al. 2022. |

| ID | Code | Type | Species | $P$ | $A$ | $S$ | $I$ | $C$ | Biome | Reference |
|---|---|---|---|---|---|---|---|---|---|---|
| 51 | sdw02b | sp | NA | 13 | 23 | 36 | 86 | 0.2876 | nt | Moracho, E et. al. 2025. |
| 52 | sdw02c | sp | NA | 6 | 9 | 15 | 19 | 0.3519 | nt | Moracho, E et. al. 2025. |
| 53 | sdw02d | sp | NA | 6 | 18 | 24 | 27 | 0.2500 | nt | Moracho, E et. al. 2025. |
| 54 | sdw03 | sp | NA | 17 | 29 | 46 | 147 | 0.2982 | nt | García-Castaño 2011. |
| 55 | sdw04 | sp | NA | 3 | 6 | 9 | 11 | 0.6111 | nt | Jordano 1993a. |
| 56 | sdw05 | sp | NA | 5 | 6 | 11 | 17 | 0.5667 | nt | Jordano 1993a. |
| 57 | sdw06 | sp | NA | 5 | 6 | 11 | 18 | 0.6000 | nt | Jordano 1993a. |
| 58 | sdw07 | sp | NA | 9 | 6 | 15 | 36 | 0.6667 | nt | Jordano 1993a. |
| 59 | sdb19 | sp | NA | 25 | 61 | 86 | 536 | 0.3515 | t | Lambert 1989. |
| 60 | sdb20 | sp | NA | 207 | 110 | 317 | 1329 | 0.0584 | t | Silva et. al. 2002. |
| 61 | sdb21 | sp | NA | 171 | 40 | 211 | 877 | 0.1282 | t | Wheelwright et. al. 1984. |
| 62 | sdw08 | sp | NA | 31 | 9 | 40 | 119 | 0.4265 | t | Beehler 1983. |
| 63 | sdw09 | sp | NA | 7 | 6 | 13 | 29 | 0.6905 | nt | Sorensen 1981. |
| 64 | sdw10 | sp | NA | 16 | 10 | 26 | 126 | 0.7875 | t | Frost 1988. |
| 65 | sdw11 | sp | NA | 12 | 7 | 19 | 52 | 0.6190 | nt | Guitian 1983. |
| 66 | sdw12 | sp | NA | 35 | 29 | 64 | 181 | 0.1783 | t | Galetti and Pizo 1996. |
| 67 | sdw13 | sp | NA | 5 | 27 | 32 | 91 | 0.6741 | t | Kantak 1981. |
| 68 | sdw14 | sp | NA | 11 | 14 | 25 | 47 | 0.3052 | nt | Snow and Snow 1988. |

| ID | Code | Type | Species | $P$ | $A$ | $S$ | $I$ | $C$ | Biome | Reference |
|---|---|---|---|---|---|---|---|---|---|---|
| 69 | sdb15 | sp | NA | 17 | 8 | 26 | 84 | 0.6176 | t | Tutin et. al. 1997. |
| 70 | sdw16 | sp | NA | 27 | 8 | 35 | 65 | 0.3009 | nt | Noma and Yumoto 1997. |
| 71 | sdw17 | sp | NA | 71 | 7 | 79 | 214 | 0.4306 | t | Crome 1975. |
| 72 | sdw18 | sp | NA | 50 | 14 | 64 | 284 | 0.4057 | t | Snow and Snow 1971. |
| 73 | sdw22 | sp | NA | 7 | 21 | 28 | 57 | 0.3878 | nt | Baird 1980. |
| 74 | sdw23 | sp | NA | 41 | 5 | 46 | 89 | 0.4341 | t | Innis 1989. |
| 75 | sdw24 | sp | NA | 15 | 49 | 64 | 158 | 0.2150 | t | Pizo 2004. |
| 76 | sdw25 | sp | NA | 22 | 13 | 35 | 123 | 0.4301 | t | Frith et. al. 1976. |
| 77 | sdb26 | sp | NA | 45 | 46 | 91 | 318 | 0.1536 | t | Donatti et. al. 2011. |
| 78 | sdb27 | sp | NA | 314 | 132 | 446 | 1902 | 0.0459 | t | Fuzessy et. al. 2022. |
| 79 | sdw28 | sp | NA | 33 | 88 | 121 | 452 | 0.1556 | t | Schleuning et. al. 2011a. |
| 80 | sdw29 | sp | NA | 49 | 16 | 65 | 180 | 0.2296 | t | Castro 2007. |
| 81 | sdw30 | sp | NA | 13 | 45 | 58 | 196 | 0.3350 | t | Correia 1997. |
| 82 | sdw31 | sp | NA | 13 | 30 | 43 | 158 | 0.4051 | t | Alves 2008. |
| 83 | sdw32 | sp | NA | 9 | 30 | 39 | 101 | 0.3741 | t | Athie 2009. |
| 84 | sdw33 | sp | NA | 25 | 28 | 53 | 115 | 0.1643 | t | Ferreira Fadini and De Marco J |
| 85 | sdw34 | sp | NA | 26 | 22 | 48 | 105 | 0.1836 | t | Hasui 1994. |
| 86 | sdw35 | sp | NA | 22 | 20 | 42 | 89 | 0.2023 | t | Silva 2011. |

| ID | Code | Type | Species | $P$ | $A$ | $S$ | $I$ | $C$ | Biome | Reference |
|---|---|---|---|---|---|---|---|---|---|---|
| 87 | sdw36-1 | sp | NA | 12 | 15 | 27 | 44 | 0.2444 | t | da Silva et. al. 2015. |
| 88 | sdw36-2 | sp | NA | 23 | 29 | 52 | 152 | 0.2279 | t | da Silva et. al. 2015. |
| 89 | sdw36-3 | sp | NA | 14 | 14 | 28 | 49 | 0.2500 | t | da Silva et. al. 2015. |
| 90 | sdw37 | sp | NA | 6 | 28 | 34 | 56 | 0.3333 | t | Robinson 2015. |
| 91 | sdw38 | sp | NA | 30 | 58 | 88 | 270 | 0.1552 | t | Rodrigues 2015. |
| 92 | sdw39 | sp | NA | 18 | 9 | 27 | 138 | 0.8519 | nt | Burns 2013. |
| 93 | sdw40 | sp | NA | 8 | 12 | 20 | 46 | 0.4792 | nt | Albrecht et. al. 2015. |
| 94 | sdw40 | sp | NA | 7 | 11 | 18 | 39 | 0.5065 | nt | Albrecht et. al. 2015. |
| 95 | sdw40 | sp | NA | 9 | 13 | 22 | 51 | 0.4359 | nt | Albrecht et. al. 2015. |
| 96 | sdw40 | sp | NA | 8 | 13 | 21 | 44 | 0.4231 | nt | Albrecht et. al. 2015. |
| 97 | sdw40 | sp | NA | 8 | 10 | 18 | 37 | 0.4625 | nt | Albrecht et. al. 2015. |
| 98 | sdw40 | sp | NA | 8 | 10 | 18 | 38 | 0.4750 | nt | Albrecht et. al. 2015. |
| 99 | sdw40 | sp | NA | 8 | 11 | 19 | 41 | 0.4659 | nt | Albrecht et. al. 2015. |
| 100 | sdw40 | sp | NA | 10 | 13 | 23 | 52 | 0.4000 | nt | Albrecht et. al. 2015. |
| 101 | sdw40 | sp | NA | 8 | 15 | 23 | 65 | 0.5417 | nt | Albrecht et. al. 2015. |
| 102 | sdw40 | sp | NA | 9 | 16 | 25 | 50 | 0.3472 | nt | Albrecht et. al. 2015. |
| 103 | sdw40 | sp | NA | 6 | 20 | 26 | 49 | 0.4083 | nt | Albrecht et. al. 2015. |
| 104 | sdw40 | sp | NA | 8 | 19 | 27 | 64 | 0.4211 | nt | Albrecht et. al. 2015. |

| ID | Code | Type | Species | $P$ | $A$ | $S$ | $I$ | $C$ | Biome | Reference |
|---|---|---|---|---|---|---|---|---|---|---|
| 105 | sdw41 | sp | NA | 22 | 17 | 39 | 100 | 0.2674 | nt | Andrade et. al. 2011. |
| 106 | sdw42 | sp | NA | 6 | 7 | 13 | 29 | 0.6905 | t | Carlo et. al. 2003. |
| 107 | sdw43 | sp | NA | 34 | 20 | 54 | 129 | 0.1897 | t | Yang et. al. 2013. |
| 108 | sdw44 | sp | NA | 14 | 6 | 20 | 36 | 0.4286 | t | Garcia et. al. 2000. |
| 109 | sdw45 | sp | NA | 81 | 18 | 99 | 282 | 0.1934 | t | Gorchov et. al. 1995. |
| 110 | sdw46 | sp | NA | 35 | 14 | 49 | 130 | 0.2653 | t | Palmeirim et. al. 1989. |
| 111 | sdw47 | sp | NA | 35 | 14 | 49 | 154 | 0.3143 | t | Lopez and Vaughan 2004. |
| 112 | sdw48 | sp | NA | 43 | 15 | 58 | 140 | 0.2171 | t | Heleno et. al. 2013. |
| 113 | sdw49 | sp | NA | 22 | 7 | 29 | 69 | 0.4481 | t | Hernandez-Montero et. al. 2015 |
| 114 | sdw49 | sp | NA | 19 | 6 | 25 | 53 | 0.4649 | t | Hernandez-Montero et. al. 2015 |
| 115 | sdw50 | sp | NA | 24 | 7 | 31 | 74 | 0.4405 | t | Passos et. al. 2003. |
| 116 | sdw51 | sp | NA | 13 | 7 | 20 | 43 | 0.4725 | t | Pedro 1992. |
| 117 | sdw52 | sp | NA | 17 | 20 | 37 | 103 | 0.3029 | t | Poulin et. al. 1999. |
| 118 | sdw53 | sp | NA | 56 | 20 | 76 | 160 | 0.1429 | t | Sarmento et. al. 2014. |
| 119 | sdw54 | sp | NA | 30 | 31 | 61 | 219 | 0.2355 | nt | Stiebel and Bairlein 2008.a. |
| 120 | sdw55 | sp | NA | 7 | 6 | 13 | 29 | 0.6905 | t | Silveira 2006. |
| 121 | sdw57 | sp | NA | 36 | 41 | 77 | 163 | 0.1104 | t | Saavedra et. al. 2014. |
| 122 | sdw57 | sp | NA | 20 | 23 | 43 | 72 | 0.1565 | t | Saavedra et. al. 2014. |

| ID | Code | Type | Species | $P$ | $A$ | $S$ | $I$ | $C$ | Biome | Reference |
|---|---|---|---|---|---|---|---|---|---|---|
| 123 | sdw58 | sp | NA | 8 | 30 | 38 | 77 | 0.3208 | nt | Menke et. al. 2012 |
| 124 | sdw58 | sp | NA | 7 | 38 | 45 | 111 | 0.4173 | nt | Menke et. al. 2012 |
| 125 | sdw58 | sp | NA | 8 | 34 | 42 | 96 | 0.3529 | nt | Menke et. al. 2012 |
| 126 | sdw58 | sp | NA | 8 | 39 | 47 | 123 | 0.3942 | nt | Menke et. al. 2012 |
| 127 | sdw59 | sp | NA | 13 | 7 | 20 | 42 | 0.4615 | t | Hayashi 1996 |
| 128 | sdw60 | sp | NA | 17 | 26 | 43 | 79 | 0.1787 | nt | Costa et. al. 2014 |
| 129 | sdw61 | sp | NA | 21 | 35 | 56 | 186 | 0.2531 | t | Giannini and Kalko 2004 |
| 130 | sdb62 | sp | NA | 787 | 343 | 1130 | 8320 | 0.0308 | t | Bello et. al. 2017 |
| 131 | sdb63 | sp | NA | 73 | 39 | 112 | 285 | 0.1001 | t | Bello et. al. 2017 |
| 133 | sdb64 | sp | NA | 47 | 42 | 89 | 163 | 0.0826 | t | Bello et. al. 2017 |
| 134 | sdb65 | sp | NA | 203 | 107 | 310 | 1124 | 0.0517 | t | Bello et. al. 2017 |
| 135 | sdb66 | sp | NA | 14 | 22 | 36 | 70 | 0.2273 | t | Kindel 1996 |
| 136 | sdw67 | sp | NA | 73 | 128 | 201 | 826 | 0.0884 | t | Stevenson et. al. 2015 |
| 137 | sdw68 | sp | NA | 14 | 42 | 56 | 119 | 0.2024 | t | Walther et. al. 2018 |
| 138 | sdb69 | sp | NA | 60 | 75 | 135 | 780 | 0.1733 | t | Palacio et. al. 2016 |
| 139 | sdw70 | sp | NA | 90 | 68 | 158 | 586 | 0.0958 | t | Machado-de Souza et. al. 2019 |
| 140 | sdw71 | sp | NA | 43 | 47 | 90 | 355 | 0.1757 | t | Naniwadekar et. al. 2019 |
| 141 | sdw72-1 | sp | NA | 140 | 98 | 238 | 572 | 0.0417 | nt | Schlautmann et. al. 2021 |

| ID | Code | Type | Species | $P$ | $A$ | $S$ | $I$ | $C$ | Biome | Reference |
|---|---|---|---|---|---|---|---|---|---|---|
| 142 | sdw72-2 | sp | NA | 7 | 14 | 21 | 25 | 0.2551 | nt | Schlautmann et. al. 2021. |
| 143 | sdw72-3 | sp | NA | 230 | 147 | 377 | 1419 | 0.0420 | nt | Schlautmann et. al. 2021. |
| 144 | sdw72-4 | sp | NA | 19 | 8 | 27 | 45 | 0.2961 | nt | Schlautmann et. al. 2021. |
| 145 | sdw72-5 | sp | NA | 153 | 133 | 286 | 795 | 0.0391 | nt | Schlautmann et. al. 2021. |
| 146 | sdw72-6 | sp | NA | 236 | 150 | 386 | 1543 | 0.0436 | nt | Schlautmann et. al. 2021. |
| 147 | sdw72-7 | sp | NA | 361 | 226 | 587 | 2398 | 0.0294 | nt | Schlautmann et. al. 2021. |
| 148 | sdw72-8 | sp | NA | 74 | 53 | 127 | 244 | 0.0622 | nt | Schlautmann et. al. 2021. |
| 149 | sdw72-9 | sp | NA | 90 | 65 | 155 | 408 | 0.0697 | nt | Schlautmann et. al. 2021. |
| 150 | sdw73 | sp | NA | 44 | 42 | 86 | 260 | 0.1407 | nt | Ramos-Robles et. al. 2016. |
| 151 | sdw74 | sp | NA | 88 | 33 | 121 | 507 | 0.1746 | t | Schleuning et. al. 2011. |
| 152 | sdw75 | sp | NA | 22 | 45 | 67 | 116 | 0.1172 | t | Dehling et. al. 2021. |
| 153 | sdw76 | sp | NA | 26 | 39 | 65 | 122 | 0.1203 | t | Dehling et. al. 2021. |
| 154 | sdw77 | sp | NA | 26 | 51 | 77 | 125 | 0.0943 | t | Dehling et. al. 2021. |
| 155 | sdw78 | sp | NA | 20 | 36 | 56 | 79 | 0.1097 | t | Dehling et. al. 2021. |
| 156 | sdw79 | sp | NA | 52 | 61 | 113 | 394 | 0.1242 | t | Dehling et. al. 2021. |
| 157 | sdw80 | sp | NA | 26 | 51 | 77 | 207 | 0.1561 | t | Dehling et. al. 2021. |
| 158 | sdw81 | sp | NA | 22 | 19 | 41 | 50 | 0.1196 | t | Dehling et. al. 2021. |
| 159 | sdw82 | sp | NA | 27 | 24 | 51 | 106 | 0.1636 | t | Dehling et. al. 2021. |

| ID | Code | Type | Species | $P$ | $A$ | $S$ | $I$ | $C$ | Biome | Reference |
|---|---|---|---|---|---|---|---|---|---|---|
| 160 | sdw83 | sp | NA | 12 | 17 | 29 | 56 | 0.2745 | nt | Dehling et. al. 2021. |
| 161 | sdw84 | sp | NA | 13 | 22 | 35 | 54 | 0.1888 | nt | Dehling et. al. 2021. |
| 162 | sdw85 | sp | NA | 14 | 25 | 39 | 55 | 0.1571 | nt | Plein et. al. 2013. |
| 163 | sdw86 | sp | NA | 13 | 24 | 37 | 67 | 0.2147 | nt | Plein et. al. 2013. |
| 164 | sdw87 | sp | NA | 18 | 26 | 44 | 73 | 0.1560 | nt | Plein et. al. 2013. |

#### Figures

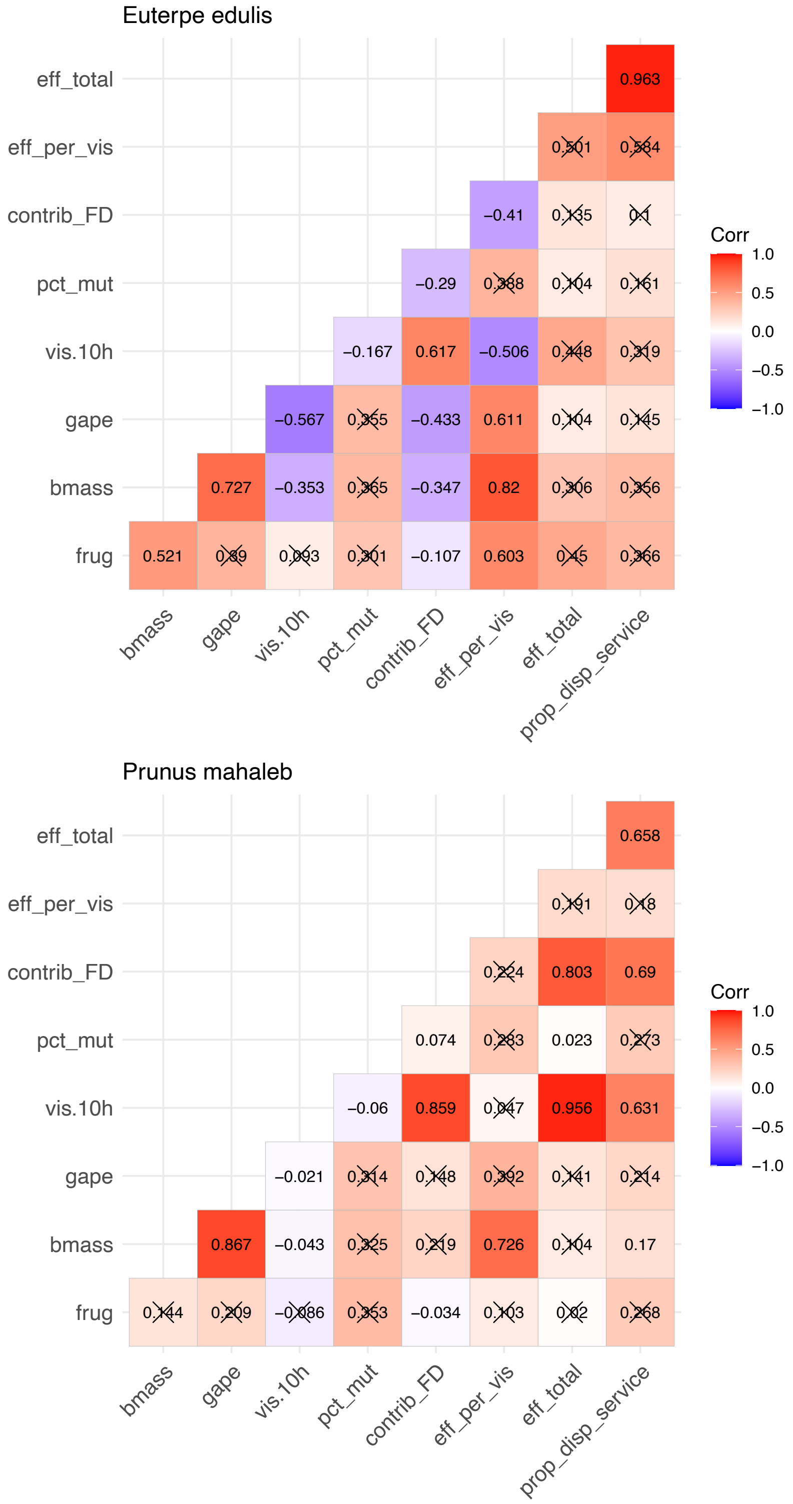


**Figure 1:** Correlation matrices among variables used to chartacterize functional diversity, $FD$. Numbers indicate the Pearson product-moment correlations between each pair of variables, with color tone proportional to the correlation value. Shown are dataresults for the two frugivore assemblages studied: *Prunus mahaleb* (Jordano and Schupp , 2000) and *Euterpe edulis* (Galetti et. al. , 2013).

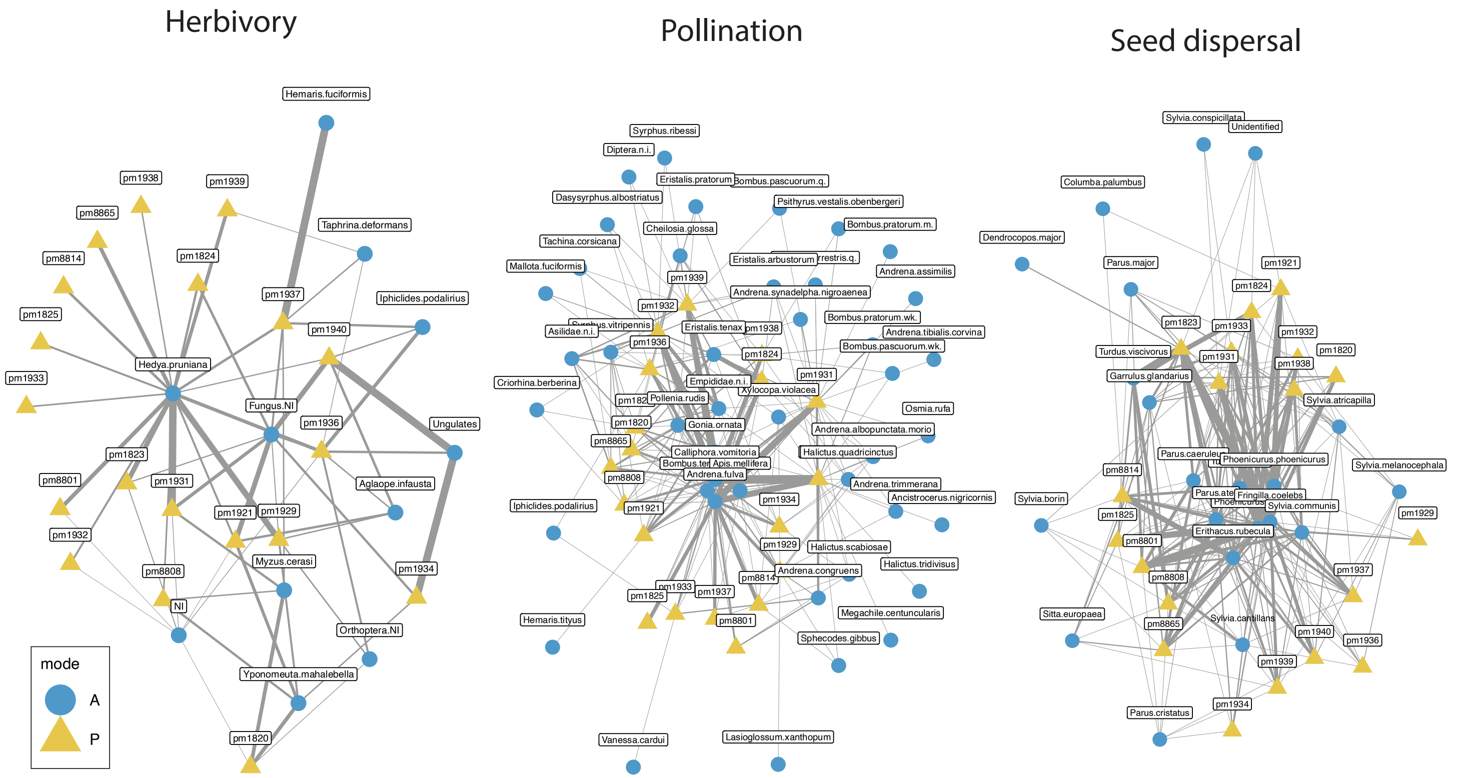


**Figure 2:** Monolayer networks of *Prunus mahaleb* tree interactions with different partners. A. Herbivory and fungi, B. Pollination, C. Seed dispersal. The width of links is proportional to the freuency of interactions between each trees and its partner taxa. Node type: yellow triangles, individual trees; blue dots, animal and fungi taxa.

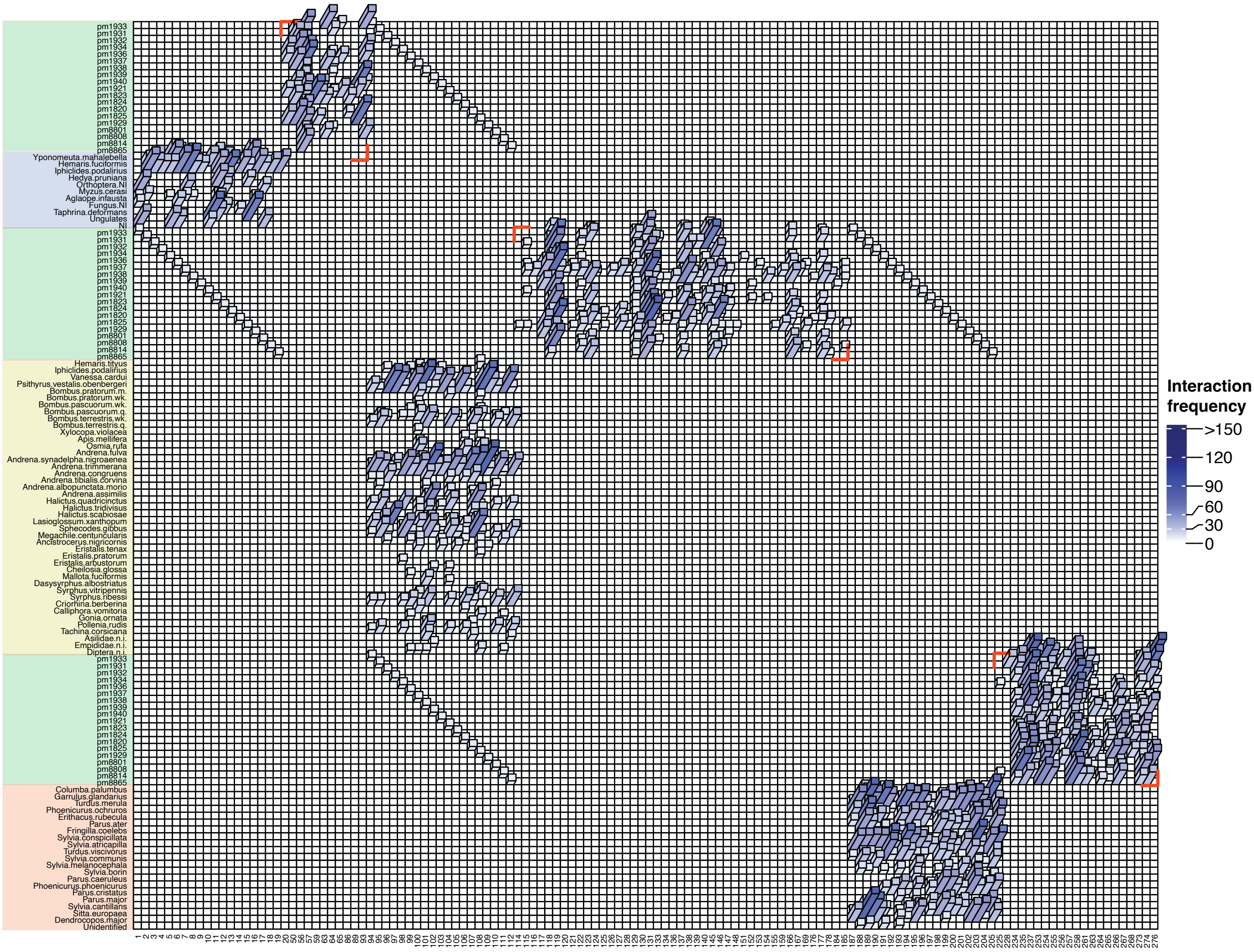


**Figure 3:** Supra-adjacency matrix of frugivore-plant interactions. The square matrix shows the number of interactions between each pair of tree (numeric labels) and animal/fungi species (scientific names). The matrix is symmetric and has the interlayer links highlighted (set to the same value to improve visualization). The color intensity is porportional to the frequency of interactions, with darker colors representing more interactions.

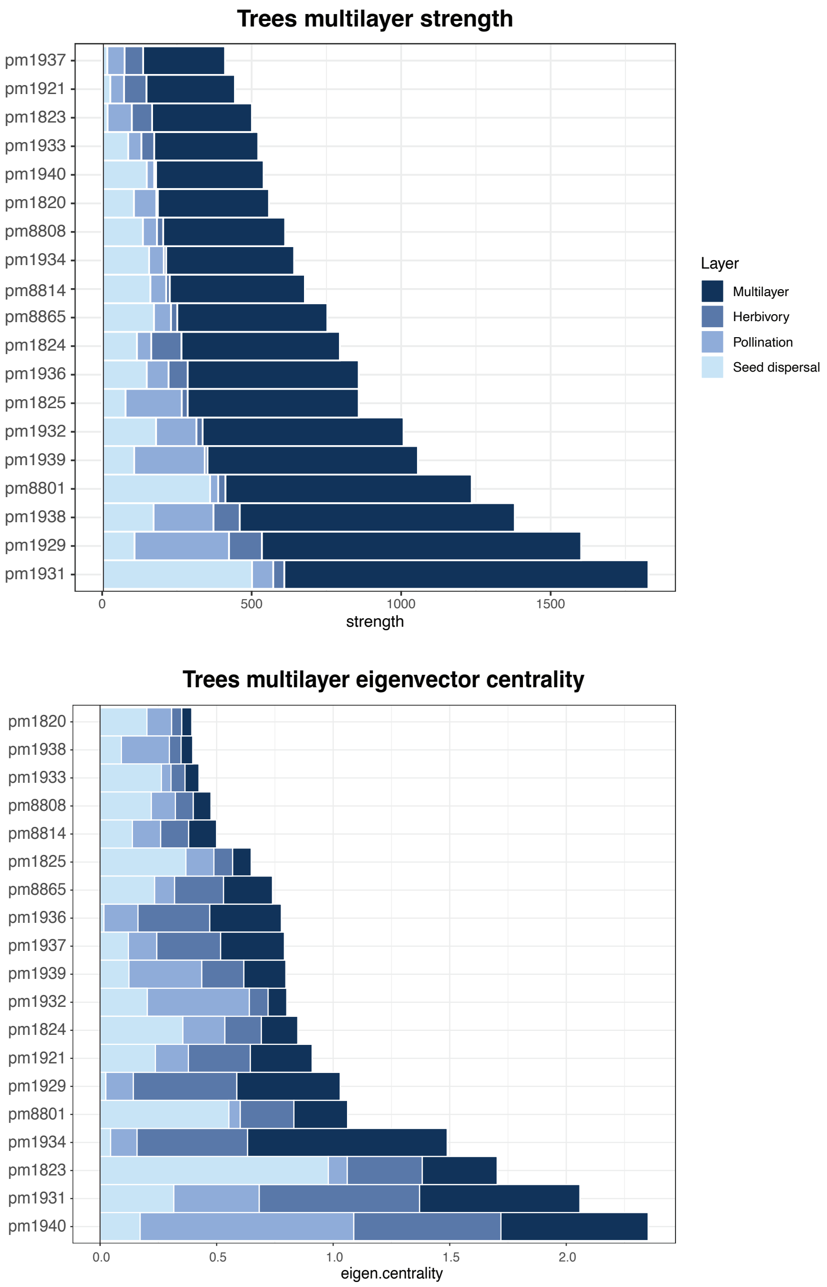


**Figure 4:** Patterns of multilayer strength (top) and multilayer centrality (bottom) for the individual *Prunus mahaleb* tress studied. The trees are ranked by their strength (top) and centrality (bottom) values in the multilayer network. Bar color indicates the type of interaction: dark blue, the whole multilayer network; lighter blue tones identify the three monolayers studied (herbivory, pollination, and seed dispersal) with decreasing blue tone.

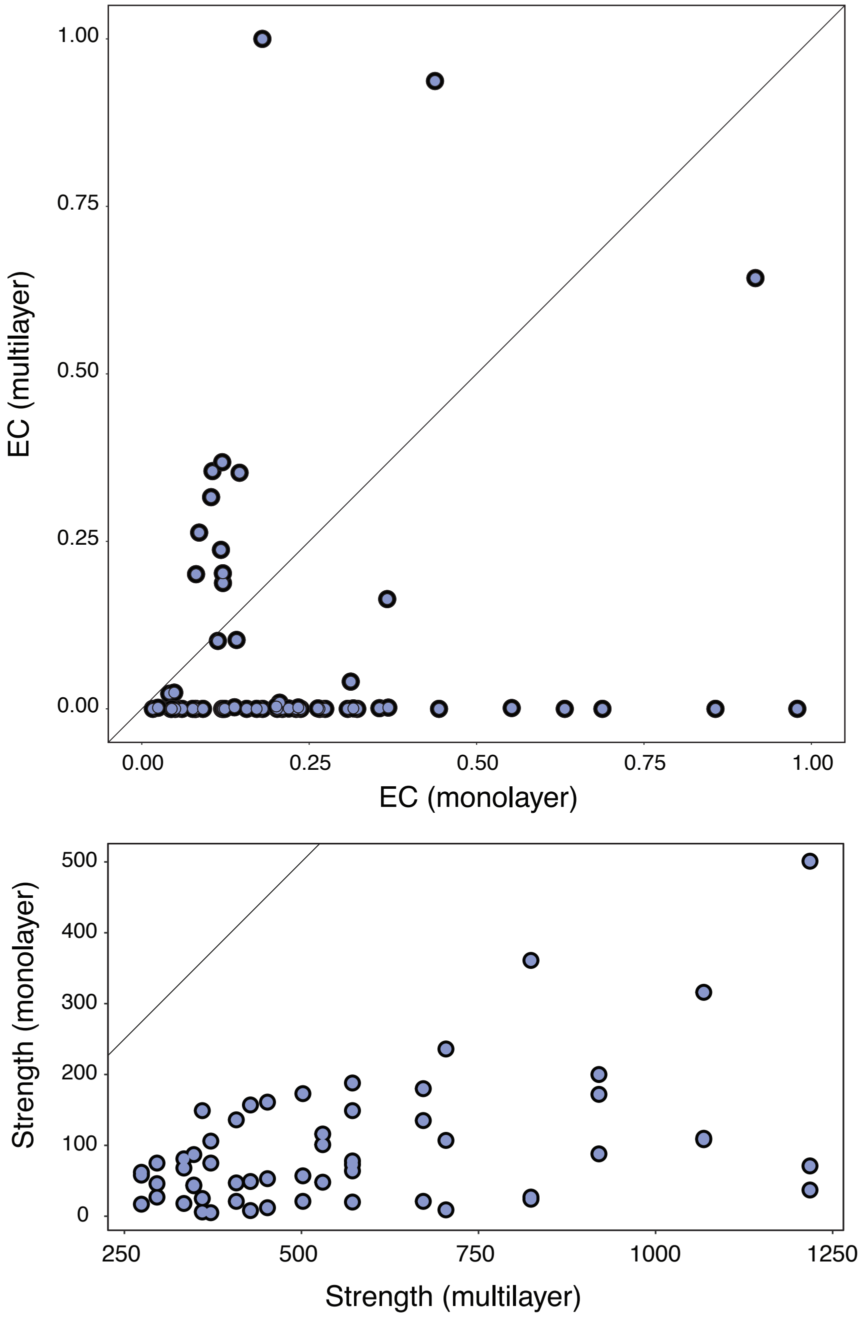


**Figure 5:** Covariation patterns between centrality (top) and strength (bottom) values for all the nodes in the network; for each node (blue dot) axes represent the values on the monolayer and the multilayer network.

Costa, J., Ramos, J. A., da Silva, L. P., Timoteo, S., Araújo, P. M., Felgueiras, M. S., Rosa, A., Matos, C., Encarnação, P., Tenreiro, P. Q. and Heleno,

R. H. 2014. Endozoochory largely outweighs epizoochory in migrating passerines. - Journal of Avian Biology 45: 59–64.

Crestani, A., Mello, M. and Cazetta, E. 2019. Interindividual variations in plant and fruit traits affect the structure of a plant-frugivore network. - Acta Oecologica 95: 120–127.

Crome, F. H. J. 1975. The ecology of fruit pigeons in tropical Northern Queensland. - Australian Wildlife Research 2: 155-185.

Dehling, D. M., Bender, I. M. A., Blendinger, P. G., Böhning-Gaese, K., Muñoz, M. C., Neuschulz, E. L., Quitián, M., Saavedra, F., Santillán, V., Schleuning, M. and Stouffer, D. B. 2021. Specialists and generalists fulfil important and complementary functional roles in ecological processes. - Functional Ecology 35: 1810–1821.

De Domenico, M., Solé-Ribalta, A., Cozzo, E., Kivelä, M., Moreno, Y., Porter, M. A., Gómez, S. and Arenas, A. 2014. Mathematical formulation of multilayer networks. - Physical Review X 3: 1–15.

De Domenico, M. 2022. Multilayer networks: analysis and visualization: introduction to muxViz with R. - Springer International Publishing.

Donatti, C. I., Guimaraes, P. R., Galetti, M., Pizo, M. A., Marquitti, F. M. D. and Dirzo, R. 2011. Analysis of a hyper-diverse seed dispersal network: modularity and underlying mechanisms. - Ecology Letters 14: 773-781.

Edler, D., Bohlin, L. and Rosvall, M. 2017. Mapping higher-order network flows in memory and multilayer networks with Infomap. - Algorithms 10: 1–23.

Ferreira Fadini, R. and De Marco Jr., P. 2004. Interacoes entre aves frugivoras e plantas em um fragmento de mata atlantica de Minas Gerais. - Ararajuba 12: 97-103.

Friedemann, P., Côrtes, M. C., Castro, E. R. d., Galetti, M., Jordano, P. and Jr, P. R. G. 2022. The individual-based network structure of palm-seed dispersers is explained by a rainforest gradient. - Oikos 2022: oik.08384.

Frith, H. J., Crome, F. H. J. and Wolfe, T. O. 1976. Food of fruit pigeons in New Guinea. - Emu 76: 49-58.

Frost, P. G. H. 1988. Fruit-frugivore interactions in a South African coastal dune forest. - In: Noring, R. (ed.), Acta XVII Congressus Internationalis Ornithologici. Deutsche Ornithologische Ges., pp. 1179-1184.

Frydman, N., Freilikhman, S., Talpaz, I. and Pilosof, S. 2023. Practical guidelines and the EMLN R package for handling ecological multilayer networks. - Methods in Ecology and Evolution 14: 2964–2973.

Fuzessy, L., Sobral, G., Carreira, D., Rother, D. C., Barbosa, G., Landis, M., Galetti, M., Dallas, T., Cláudio, V. C., Culot, L. and Jordano, P. 2022. Functional roles of frugivores and plants shape hyper-diverse mutualistic interactions under two antagonistic conservation scenarios. - Biotropica 54: 444–454.

Galetti, M., Guevara, R., Côrtes, M. C., Fadini, R., Von Matter, S., Leite, A. B., Labecca, F., Ribeiro, T., Carvalho, C. S., Collevatti, R. G., Pires, M. M., Guimarães Jr., P. R., Brancalion, P. H., Ribeiro, M. C. and Jordano, P. 2013. Functional extinction of birds drives rapid evolutionary changes in seed size. - Science 340: 1086–1091.

Galetti, M. and Pizo, M. A. 1996. Fruit eating birds in a forest fragment in southeastern Brazil. - Ararajuba, Rev. Brasil. Ornitol. 4: 71-79.

Garcia, Q. S., Rezende, J. L. P. and Aguiar, L. M. S. 2000. Seed dispersal by bats in a disturbed area of Southeastern Brazil. - Revista de Biología Tropical 48: 125-128.

García-Castaño, J. L. 2011. Consecuencias demográficas de la dispersión de semillas por aves y mamíferos frugívoros en la vegetación mediterránea de montaña. - Universidad de Sevilla.

Giannini, N. P. and Kalko, E. K. 2004. Trophic structure in a large assemblage of phyllostomid bats in Panama. - Oikos 105: 209–220.

Gopal, A., Mudappa, D., Raman, T. and Naniwadekar, R. 2020. Forest cover and fruit crop size differentially influence frugivory of select rainforest tree species in Western Ghats, India. - Biotropica 52: 871 - 883.

Gorchov, D. L., Cornejo, F., Ascorra, C. F. and Jaramillo, M. 1995. Dietary overlap between frugivorous birds and bats in the peruvian amazon. - Oikos 74: 235-250.

Guerra, T. J., Dayrell, R. L. C., Arruda, A. J., Dáttilo, W., Teixido, A. L., Messeder, J. V. S. and Silveira, F. A. O. 2017. Intraspecific variation in fruit–frugivore interactions: effects of fruiting neighborhood and consequences for seed dispersal. - Oecologia 185: 233–243.

Guitian, J. 1983. Relaciones entre los frutos y los passeriformes en un bosque montano de la cordillera cantábrica occidental. - Universidad de Santiago, Spain.

Hasui, E. 1994. O papel das aves frugívoras na dispersão de sementes em um fragmento de floresta semidecídua secundaria em São Paulo, SP. - Universidade de São Paulo.

Hayashi, M. M. 1996. Morcegos frugívoros em duas áreas alteradas da fazenda Lageado, Botucatu, Estado de São Paulo. - Universidade Estadual Paulista.

Heleno, R. H., Olesen, J. M., Nogales, M., Vargas, P. and Traveset, A. 2013. Seed dispersal networks in the Galapagos and the consequences of alien plant invasions. - Proceedings of the Royal Society, Biol. Sci. 280: 20122112.

Hernandez-Montero, J. R., Saldana-Vazquez, R. A., Galindo-Gonzalez, J. and Sosa, V. J. 2015. Bat-fruit interactions are more specialized in shaded-coffee plantations than in tropical mountain cloud forest fragments. - PLoS One 10: e0126084.

Hutchinson, M. C., Bramon Mora, B., Pilosof, S., Barner, A. K., Kéfi, S., Thébault, E., Jordano, P. and Stouffer, D. B. 2019. Seeing the forest for the trees: Putting multilayer networks to work for community ecology. - Functional Ecology 33: 206–217.

Innis, G. J. 1989. Feeding ecology of fruit pigeons in subtropical rainforests of south-eastern Queensland. - Australian Journal of Wildlife Research 16: 365-394.

Isla, J., Jácome-Flores, M., Arroyo, J. M. and Jordano, P. 2023. The turnover of plant–frugivore interactions along plant range expansion: consequences for natural colonization processes. - Proceedings of the Royal Society B: Biological Sciences 290: 20222547.

Jayanth, A., Isvaran, K. and Naniwadekar, R. 2024. Drivers of intraspecific variation in seed dispersal can differ across two species of fleshy-fruited savanna plants. - Biotropica 56: e13322.

Jordano, P. 1993. Pollination biology of Prunus mahaleb L.: deferred consequences of gender variation for fecundity and seed size. - Biological Journal of the Linnean Society 50: 65–84.

Jordano, P. 1993a. Geographical ecology and variation of plant-seed disperser interactions: southern Spanish junipers and frugivorous thrushes. - In: Fleming, T. H. and Estrada, A. (ed.), Frugivory and seed dispersal: ecological and evolutionary aspects. Kluwer Academic Publisher, pp. 85–104.

Jordano, P. 1995. Frugivore-mediated selection on fruit and seed size: birds and St. Lucie's cherry, Prunus mahaleb. - Ecology 76: 2627–2627.

Jordano, P. and Schupp, E. W. 2000. Seed disperser effectiveness: The quantity component and patterns of seed rain for *Prunus mahaleb*. - Ecological Monographs 70: 591–615.

Jácome-Flores, M. E., Jordano, P., Delibes, M. and Fedriani, J. M. 2020. Interaction motifs variability in a Mediterranean palm under environmental disturbances: the mutualism–antagonism continuum. - Oikos 129: 367–379.

Kantak, G. E. 1981. Temporal feeding patterns of some tropical frugivores. - Condor 83: 185-187.

Kindel, A. 1996. Interações entre plantas ornitocóricas e aves frugívoras na Estação Ecológica de Aracuri, Muitos Capões, RS. - Universidade Federal de Rio Grande do Sul.

Kivelä, M., Arenas, A., Barthelemy, M., Gleeson, J. P., Moreno, Y., Porter, M. A., Kivelå, M., Arenas, A., Barthelemy, M., Gleeson, J. P., Moreno, Y.,

Porter, M. A., Kivelä, M., Arenas, A., Barthelemy, M., Gleeson, J. P., Moreno, Y. and Porter, M. A. 2014. Multilayer networks. - Journal of Complex Networks 2: 203–271.

Lambert, F. 1989. Fig-eating by birds in a Malaysian lowland rain forest. - Journal of Tropical Ecology 5: 401-412.

Lamperty, T., Karubian, J. and Dunham, A. E. 2021. Ecological drivers of intraspecific variation in seed dispersal services of a common neotropical palm. - Biotropica 53: 1226–1237.

Lopez, J. E. and Vaughan, C. 2004. Observations on the Role of Frugivorous Bats as Seed Dispersers in Costa Rican Secondary Humid Forests. - Acta Chiropterologica 6: 111-119.

Machado-de Souza, T., Campos, R. P., Devoto, M. and Varassin, I. G. 2019. Local drivers of the structure of a tropical bird-seed dispersal network. - Oecologia 189: 421–433.

Menke, S., Böhning-Gaese, K. and Schleuning, M. 2012. Plant-frugivore networks are less specialized and more robust at forest-farmland edges than in the interior of a tropical forest. - Oikos 121: 1553–1566.

Miguel, M. F., Jordano, P., Tabeni, S. and Campos, C. M. 2018. Context-dependency and anthropogenic effects on individual plant–frugivore networks. - Oikos 127: 1045–1059.

Naniwadekar, R., Chaplod, S., Datta, A., Rathore, A. and Sridhar, H. 2019. Large frugivores matter: Insights from network and seed dispersal effectiveness approaches. - Journal of Animal Ecology 88: 1250–1262.

Noma, N. and Yumoto, T. 1997. Fruiting phenology of animal-dispersed plants in response to winter migration of frugivores in a warm temperate forest on Yakushima Island, Japan. - Ecological Research 12: 119-129.

Palacio, R. D., Valderrama-Ardila, C. and Kattan, G. H. 2016. Generalist species have a central role in a highly diverse plant–frugivore network. - Biotropica 48: 349–355.

Palmeirim, J. M., Gorchov, D. L. and Stoleson, S. 1989. Trophic structure of a neotropical frugivore community: is there competition between birds and bats? - Oecologia (Berl.) 79: 403-411.

Passos, F. C., Silva, W. R., Pedro, W. A. and Bonin, M. R. 2003. Frugivoria em morcegos (Mammalia, Chiroptera) no Parque Estadual Intervales, sudeste do Brasil. - Revista Brasileira de Zoologia 20: 511-517.

Pedro, W. 1992. Estrutura de uma taxocenose de morcegos da Reserva do Panga (Uberlândia, MG), com ênfase nas relações tróficas em Phyllostomidae (Mammalia: Chiroptera). - Universidade Estadual de Campinas.

Pilosof, S., Porter, M. A., Pascual, M. and Kéfi, S. 2017. The multilayer nature of ecological networks. - Nature Ecology & Evolution 1: 0101.

Pizo, M. A. 2004. Frugivory and habitat use by fruit-eating birds in a fragmented landscape of southeast Brazil. - Ornitologia Neotropical 15 Suppl.: 117–126.

Plein, M., Längsfeld, L., Neuschulz, E. L., Schultheib, C., Ingmann, L., Töpfer, T., Böhning-Gaese, K. and Schleuning, M. 2013. Constant properties of plant-frugivore networks despite fluctuations in fruit and bird communities in space and time. - Ecology 94: 1296–1306.

Poulin, B., Wright, S. J., Lefebvre, G. and Calderon, O. 1999. Interspecific synchrony and asynchrony in the fruiting phenologies of congeneric bird-dispersed plants in Panama. - Journal of Tropical Ecology 15: 213-227.

Quintero, E., Isla, J. and Jordano, P. 2022. Methodological overview and data-merging approaches in the study of plant–frugivore interactions. - Oikos : e08379.

Quintero, E., Rodríguez-Sánchez, F. and Jordano, P. 2023. Reciprocity and interaction effectiveness in generalised mutualisms among free-living species. - Ecology Letters 26: 132–146.

Ramaswami, G., Somnath, P. and Quader, S. 2017. Plant-disperser mutualisms in a semi-arid habitat invaded by Lantana camara L. - Plant Ecology 218: 935–946.

Ramos-Robles, M., Andresen, E. and Díaz-Castelazo, C. 2016. Temporal changes in the structure of a plant-frugivore network are influenced by bird migration and fruit availability. - PeerJ 4: e2048–e2048.

Robinson, V. 2015. Índice de importancia de diferentes especies de plantas na atração de aves para uma area reflorestada em Piracicaba. - Universidade Estadual Paulista "Júlio de Mesquita Filho".

Rodrigues, S. 2015. Rede de interaçoes entre aves frugívoras e plantas em uma area de Mata Atlântica no sudeste do Brasil. - Universidade Federal de Sâo Carlos, Campus Sorocaba.

Saavedra, F., Hensen, I., Beck, S. G., Böhning-Gaese, K., Lippok, D., Töpfer, T. and Schleuning, M. 2014. Functional importance of avian seed dispersers changes in response to human-induced forest edges in tropical seed-dispersal networks. - Oecologia 176: 837–848.

Sarmento, R., Alves-Costa, C. P., Ayub, A. and Mello, M. A. R. 2014. Partitioning of seed dispersal services between birds and bats in a fragment of the Brazilian Atlantic Forest. - Zoologia (Curitiba, Impresso) 31: 245-255.

Schlautmann, J., Rehling, F., Albrecht, J., Jaroszewicz, B., Schabo, D. G. and Farwig, N. 2021. Observing frugivores or collecting scats: a method comparison to construct quantitative seed dispersal networks. - Oikos 130: 1359–1369.

Schleuning, M., Blüthgen, N., Florchinger, M., Braun, J., Schaefer, H. M. and Böhning-Gaese, K. 2011. Specialization and interaction strength in a tropical plant-frugivore network differ among forest strata. - Ecology 92: 26–36.

Schleuning, M., Bluthgen, N., Florchinger, M., Braun, J., Schaefer, H. M. and Bohning-Gaese, K. 2011a. Specialization and interaction strength in a tropical plant-frugivore network differ among forest strata. - Ecology 92: 26-36.

Silva, R. d. M. 2011. Interações entre plantas e aves frugívoras no campus da Universidade Federal do Rio de Janeiro. - Universidade Federal do Rio de Janeiro.

Silva, W. R., De Marco Jr, P., Hasui, E. and Gomes, V. S. M. 2002. Patterns of fruit-frugivore interactions in two atlantic forest bird communities of South-eastern Brazil: implications for conservation. - In: Levey, D. J., Silva, W. R. and Galetti, M. (ed.), Seed dispersal and frugivory: ecology, evolution and conservation. CAB International, pp. 423-435.

Silveira, M. 2006. Dispersão de sementes por morcegos frugívoros em uma área em processo de restauração vegetal na RPPN Parque Florestal São Marcelo Mogi-Guaçu, SP. - Universidade Estadual de São Paulo.

Snow, B. K. and Snow, D. W. 1971. The feeding ecology of tanagers and honeycreepers in Trinidad. - Auk 88: 291-322.

Snow, B. K. and Snow, D. W. 1988. Birds and berries. - Poyser, Calton, UK.

Sorensen, A. E. 1981. Interactions between birds and fruit in a temperate woodland. - Oecologia (Berl.) 50: 242-249.

Stevenson, P. R., Link, A., González-Caro, S. and Torres-Jiménez, M. F. 2015. Frugivory in canopy plants in a western Amazonian forest: dispersal systems, phylogenetic ensembles and keystone plants. - PLoS One 10: e0140751–22.

Stiebel, H. and Bairlein, F. 2008. Frugivorie mitteleuropäischer Vögel I: Nahrung und Nahrungserwerb. - Vogelwarte 46: 1-23.

Stiebel, H. and Bairlein, F. 2008a. Frugivorie mitteleuropäischer Vögel II: Einfluss des Fruchtangebotes auf die raäumliche und zeitliche Habitatnutzung frugivorer Vogelarten. - Vogelwarte 46: 81-94.

Thiel, S., Gottstein, M., Heymann, E. W., Kroszewski, J., Lieker, N., Tello, N. S., Tschapka, M., Junker, R. R. and Heer, K. 2024. Vertically stratified interactions of nectarivores and nectar-inhabiting bacteria in a liana flowering across forest strata*. - American Journal of Botany 111: e16303.

Tutin, C. E. G., Ham, R. M., White, L. J. T. and Harrison, M. J. S. 1997. The primate community of the Lopé Reserve, Gabon: diets, responses to fruit scarcity, and effects on biomass. - Am. J. Primatol. 42: 1-24.

Vergara-Tabares, D. L., Blendinger, P. G., Tello, A., Peluc, S. I. and Tecco, P. A. 2022. Fleshy-fruited invasive shrubs indirectly increase native tree seed dispersal. - Oikos 2022: e08311.

Villalva, P., Arroyo-Correa, B., Calvo, G., Homet, P., Isla, J., Mendoza, I., Moracho, E., Quintero, E., Rodríguez-Sánchez, F. and Jordano, P. 2024.

FRUGIVORY CAMTRAP: A dataset of plant–animal interactions recorded with camera traps. - Ecology 105: e4424.

Vissoto, M., Vizentin-Bugoni, J., Sendoya, S. F., Gomes, G. C. and Dias, R. A. 2022. Plant height and spatial context influence individual connectivity and specialization on seed dispersers in a tree population. - Oecologia 198: 721–731.

Walther, B. A., Geier, J., Chou, L.-S. and Bain, A. 2018. The figs of winter: Seasonal importance of fruiting fig trees (Ficus: Moraceae) for urban birds. - Acta Oecologica 90: 28–34.

Wheelwright, N. T., Haber, W. A., Murray, K. G. and Guindon, C. 1984. Tropical fruit-eating birds and their food plants: a survey of a Costa Rican lower montane forest. - Biotropica 16: 173-192.

Yang, S., Albert, R. and Carlo, T. A. 2013. Transience and constancy of interactions in a plant-frugivore network. - Ecosphere 4: art147.

da Silva, F., Montoya, D., Furtado, R., Memmott, J., Pizo, M. A. and Rodrigues, R. R. 2015. The restoration of tropical seed dispersal networks. - Restoration Ecology 23: 852-860.

Jordano, P., 2007. A compilation of plant-frugivore interactions for Spanish habitats. v.1.0.0. - : .

Jordano, P., 2024. La biodiversidad de las interacciones ecológicas. - In: Medel, R., , Traveset, A., and Navarro, L., (ed.), Más allá de la pérdida de especies. Interacciones ecológicas en el Antropoceno. Ediciones Fremen SpA, pp. 16–49.

Jordano, P., 2024a. Material for the course: Curso Posgraduação Frugivoria 2016, UNESP, Rio Claro, Brazil. doi: 10.5281/zenodo.10478535. - : .

Moracho, E, , Arroyo-Correa, B., , Calvo, G., , Homet, P., , Isla, J., , Mendoza, I., , Quintero, E., , Villalva, P., , Rodríguez-Sánchez, F., and Jordano, P., 2025. DOÑANA-FRUGINT: A dataset of plant-animal interactions. - MS 00: 000–000.

Olesen, JM, , Bascompte, J., Elberling, H. and Jordano, P. 2008. Temporal dynamics in a pollination network. - Ecology 89: 1573–1582.

Rodríguez-Sánchez, F., 2010. An integrative framework to investigate species responses to climate change: Biogeography and ecology of relict trees in the Mediterranean. Ph.D. Thesis. - Universidad de Sevilla.
